# Characterization of a novel mouse model of Dopamine Transporter Deficiency Syndrome and pharmacological therapeutic strategies

**DOI:** 10.1101/2025.03.21.644417

**Authors:** Emma E. Russo, Ameneh Rezayof, Conner Wallace, Erin Q. Williams, Pieter Beerepoot, Marija Milenkovic, Maria Novalen, Aled Blundell, Tatiana V. Lipina, Jason Locke, Raveen Christian, Peter S.B Finnie, Landon J. Edgar, Rachel F. Tyndale, Dawn Watkins-Chow, Amy J. Ramsey, Sara R. Jones, Ali Salahpour

## Abstract

The dopamine transporter (DAT) is an essential protein in the maintenance of dopamine homeostasis in the brain. Thus, single amino acid changes in the gene that encodes for DAT can be sufficient to induce disease, such as Dopamine Transporter Deficiency Syndrome (DTDS). DTDS-associated variants are posited to cause DAT protein misfolding, retention in the endoplasmic reticulum, and a consequent depletion or loss of DAT at the cell surface. In turn, proper dopaminergic regulation is lost. Current treatments for DTDS are largely ineffective, and improved therapeutic options are greatly needed. To this end, we have created a novel mouse model of DTDS harboring the A313V knock-in DAT variant, a proxy for the DTDS-causing A314V variant in humans. We show that the A313V knock-in DAT mice are hyperactive, have increased striatal tissue content of dopamine and its metabolites homovanillic acid (HVA) and DOPAC, and impaired dopamine uptake. We demonstrate that FDA approved compounds alpha-methyl-para-tyrosine (ɑMPT) and amphetamine (AMPH) ameliorate hyperactivity in this mouse model. Moreover, ɑMPT may be a disease-modifying treatment by addressing the hyperdopaminergic tone underlying this hyperactivity. In contrast, noribogaine, a pharmacological chaperone for DAT, is unable to rescue DAT expression. Taken together, these findings show that the A313V knock-in DAT variant mice recapitulate several defining phenotypes seen in patients with DTDS, and provide evidence for two novel treatments for the disease.

## Introduction

The dopamine transporter (DAT) is a twelve transmembrane domain protein that is expressed in dopamine neurons (1–4). DAT is critically involved in dopamine (DA) homeostasis by coupling the transport of extracellular DA with Na+ and Cl-back into the presynaptic neuron (5–7), similar to other members of the neurotransmitter symporter solute carrier 6 (SLC6) family, including the norepinephrine transporter, the serotonin transporter, and the GABA transporter (8).

Variants in the DAT gene, SLC6A3, have been implicated in several diseases (9) including Autism Spectrum Disorder (10,11), Attention Deficit Hyperactivity Disorder (ADHD) (12), Bipolar Disorder (13), and Dopamine Transporter Deficiency Syndrome (DTDS) (14–26). DTDS is typically characterized first by symptoms of hyperactivity; these symptoms progress over time to a state of parkinsonism-dystonia through unknown mechanisms and are accompanied by increased levels of DA metabolites present in cerebrospinal fluid (Ng et al., 2014). Patients diagnosed with DTDS rarely live beyond adolescence due to secondary complications of the disease, such as orthopedic, gastrointestinal, and respiratory issues (16, 17, 27). Importantly, DTDS is considered a “pharmaco-resistant condition” (17). Treatment strategies such as muscle relaxants, dopaminergic, anticholinergic, and GABAergic medications, along with surgical interventions, have generally proven unsuccessful (15–17). Thus, effective treatments for DTDS are greatly needed.

SLC6A3 missense variants associated with DTDS are posited to cause protein misfolding, which disrupts trafficking of DAT and reduces protein levels at the membrane (15–17) needed to maintain typical physiological functions (28). The quality control machinery in the endoplasmic reticulum (ER) normally prevents misfolded proteins with aberrant conformations from advancing along the secretory pathway. However, the ER can excessively retain and degrade proteins that are potentially functional, leading to inadequate levels of the protein at its physiological destination (29). In the case of DTDS-causing variants, impeded DAT maturation through the ER leads to low levels or no surface expression of the protein, reduced DA uptake, and consequent disease (15–17, 30, 31).

Pharmacological chaperones are exogenously-applied, small molecule ligands that bind to an intermediately-folded protein of interest and assist in its exit from the ER, providing an opportunity for “rescue” of misfolded membrane proteins. This “rescue” is proposed to be mediated through stabilization of a native-like conformation of the target protein, allowing it to exit from the ER (32, 33). Successful rescue with pharmacological chaperones has been demonstrated for clinically-relevant misfolded disease-causing variants of several proteins, such as the V2 vasopressin receptor (34), alpha-galactosidase A (35), the F508del cystic fibrosis transmembrane conductance receptor (36) and the gonadotropin releasing hormone receptor (37). SLC6 transporters are likewise amenable to rescue via pharmacological chaperones (38), as demonstrated for SERT (33, 39, 40) and DAT (30, 31, 33, 41). For both SERT and DAT, rescue can be achieved using ibogaine and noribogaine, the less cardiotoxic (42) active metabolite of ibogaine, as well as bupropion (30, 31, 33, 41, 43).

In order to gain further insight into DTDS and identify potential pharmacotherapies, we have generated a novel line of transgenic knock-in mice carrying one of the DTDS missense variants. The knock-in mouse (mA313V variant) models the human DTDS hA314V variant and was chosen for this study because 1.) patients harboring the hA314V variant can live to adulthood (17), providing a window for pharmacological intervention, and 2.) this hA314V DAT variant can be rescued by pharmacological chaperones *in vitro* (30). In this study, we show that the mA313V mice recapitulate a subset of defining phenotypes observed in human DTDS patients, including hyperlocomotion and changes in DA metabolites. The mA313V DAT model also has reduced expression and uptake of DA, providing the first support from a mammalian model for observations originally made in heterologous cell systems. Our attempts to rescue the ER-retained, misfolded A313V DTDS variant using the pharmacological chaperone noribogaine were unsuccessful. However, findings show that AMPH and the tyrosine hydroxylase inhibitor alpha-methyl-para-tyrosine (ɑMPT) can effectively reduce the hyperactivity of the DTDS animals. These key findings are of both clinical and translational interest. Indeed, ɑMPT, an FDA-approved drug that is used for clinical management of hypertension, 1.) reduces hyperactivity of the mA313V mice, and 2.) may have disease modifying properties and reduce disease burden by inhibiting excess dopamine synthesis and neurotransmission. Altogether, our study reports upon the efficacy of pharmacological treatment via clinically available drugs that may be translated to human DTDS patients.

## Results

### Neurochemical Characterization

LLC-PK_1_ and HEK cells harboring the DTDS-causing human A314V SLC6A3 variant were reported to express reduced mature (fully-glycosylated) DAT and an increased ratio of immature to mature (core-glycosylated) DAT protein (17, 30). However, to date, no study has assessed the maturation status of a DTDS-causing variant in a mammalian model. As shown in Fig. 1*a-d*, striatal (100 ± 9.93 vs 23.01± 5.79 % of WT average, t=6.115, df=8, p=0.0003) and midbrain (100 ± 6.77 vs 24.39 ± 3.57 % of WT average, t=9.876, df=6, p<0.0001) homogenates from the A313V knock-in mice express significantly decreased levels of mature DAT (Fig. 1*a-d*) and an increased percentage of immature DAT to total DAT levels in the midbrain (3.66 ± 0.20 vs 31.67 ± 3.73 % immature DAT, t=7.499, df=6, p=0.0003) (Fig. 1*e*). Indeed, there is a 75% decrease in mature DAT protein expression in the striatum and midbrain of A313V mice as compared to WT littermate controls, while there is a marked (∼6 fold) increase in immature protein in the midbrains of A313V mice. To confirm that the higher molecular weight band is mature DAT, while the lower molecular weight band is immature DAT, midbrain homogenates were next digested with the glycosidases peptide *N*-glycosidase F (PNGase F) and endoglycosidase H (EndoH). As shown in Fig 2, results confirm that the upper molecular weight band in the brain western blot samples indeed represents the mature, fully-glycosylated DAT protein, whereas the lower band represents the immature, core-glycosylated DAT protein localized to the ER (Fig. 2). This is the first evidence that a DTDS-disease causing variant leads to DAT retention in the ER and an increase in immature DAT protein species in a mammalian model.

**Figure 1.**
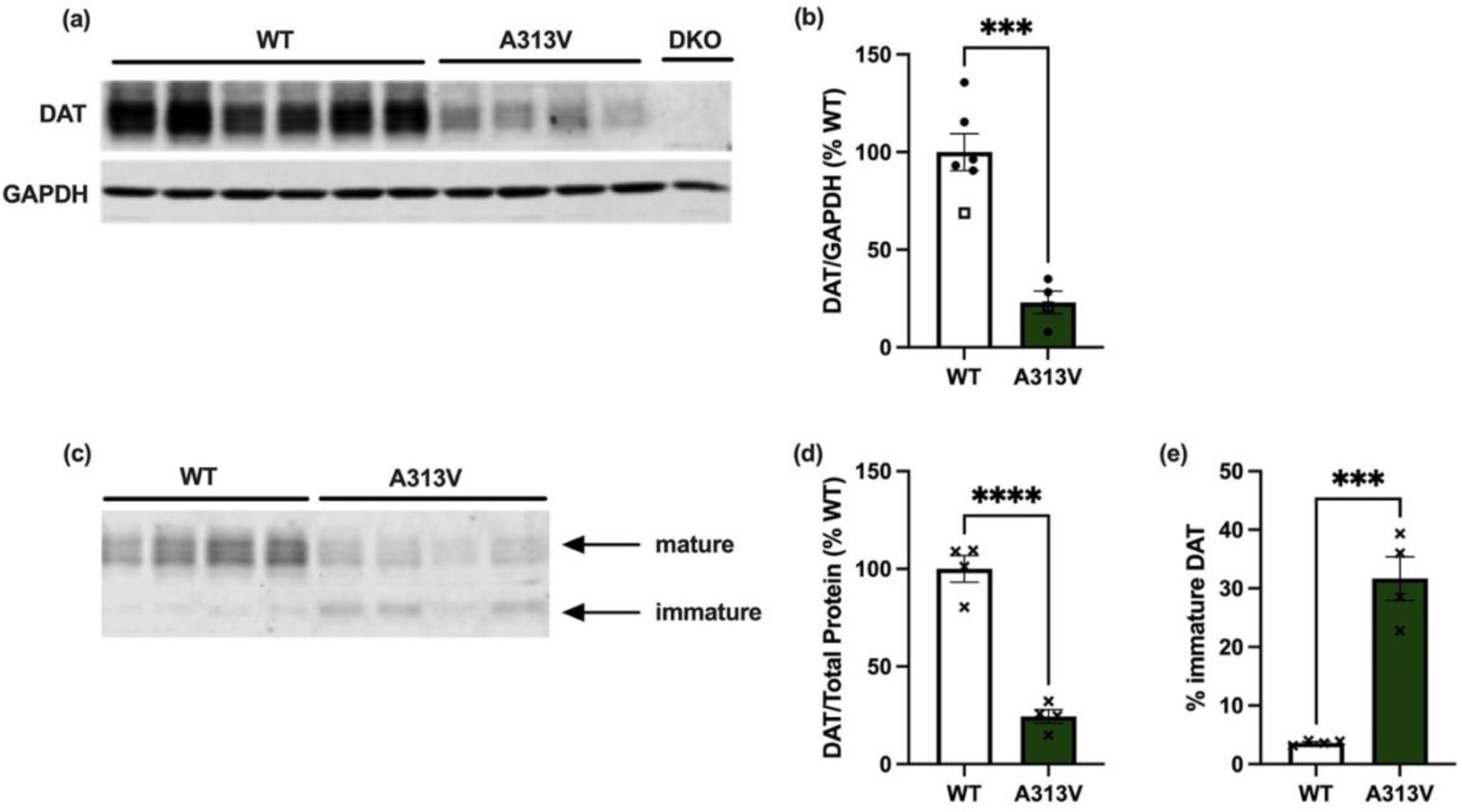
Western blot analysis of striatal and midbrain DAT levels in WT and A313V mice. (a) Representative western blot depicting striatal dopamine transporter (DAT) levels in wildtype (WT) and DAT A313V-knock in (A313V) mice, with a DAT knock-out (DKO) negative control (n=4-6). (b) Quantification of striatal DAT levels in WT and A313V mice relative to WT levels and normalized to GAPDH loading control showing A313V have significantly less striatal DAT compared to WT mice (100 ± 9.93 vs 23.01± 5.79 % of WT average, t=6.115, df=8, p=0.0003). (c) Representative western blot depicting DAT levels in WT and A313V. Arrows indicate the mature and immature forms of the DAT protein (n=4, 3 midbrains/lane). (d) Quantification of midbrain DAT levels in WT and A313V mice relative to WT levels and normalized to total protein loading control showing A313V mice have significantly lower levels of midbrain DAT compared to WT mice (100 ± 6.77 vs 24.39 ± 3.57 % of WT average, t=9.876, df=6, p<0.0001). (e) Bar graph quantification showing the percentage of immature DAT levels to total DAT levels in the midbrain in WT and A313V mice demonstrating A313V mice have a greater percentage of immature DAT respective to total DAT compared to WT mice (3.66 ± 0.20 vs 31.67 ± 3.73 % immature DAT, t=7.499, df=6, p=0.0003). Closed circular points represent values from female mice, and open square points represent values from male mice. X’s represent value from mixed-sex samples. Student’s unpaired, two-tailed *t-*tests were conducted. All results are presented as mean ± SEM, ***p≤0.001, ****p<0.0001

**Figure 2.**
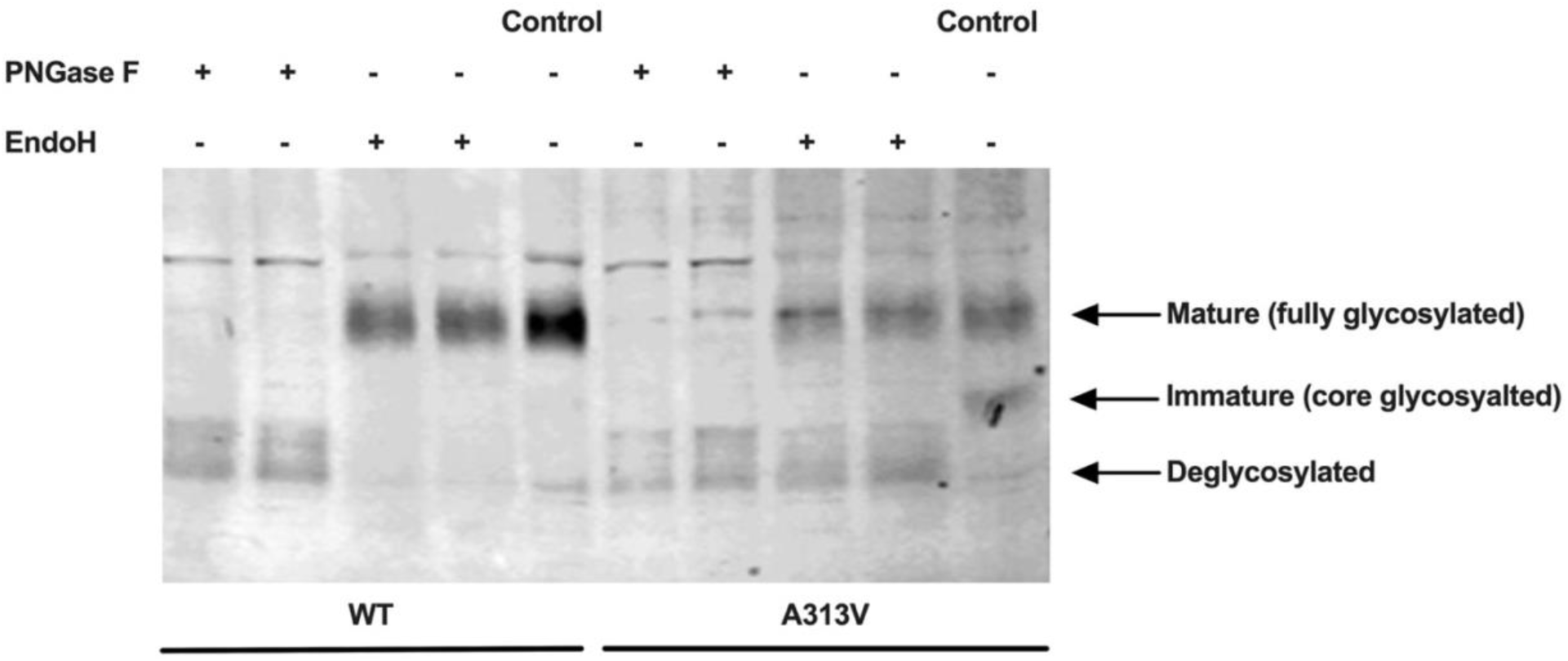
Analysis of Midbrain DAT Protein Species in WT and A313V Mice. The higher molecular weight band represents the mature, fully glycosylated dopamine transporter (DAT) with complex oligosaccharides, which were removed with the enzyme PNGase F in wildtype (WT) (lanes 1 and 2) and DAT A313V knock-in (A313V) (lanes 6 and 7) mice. The lower molecular weight band represents the immature, core glycosylated DAT as evident by digestion with the enzyme endoglycosidase (EndoH) in A313V midbrain samples (lanes 8 and 9). Each lane has 3 midbrains per sample, with mixed-sexes in each sample.

Lower abundance of mature DAT in the striatum suggests that A313V mice may have impaired presynaptic DA recycling, and thus depleted reserve pools of dopamine, similar to what has been described in DAT-knock out (DAT-KO) animals (44, 45). Analysis of striatal tissue content of dopamine and its metabolites revealed that the A313V mice have a dramatic (approximately four-fold) decrease in dopamine tissue levels (471.23 ± 47.51 vs 104.13 ± 6.35 pg/mg, t=10.63, df=9, p<0.0001) and nearly 50% less DOPAC compared to WT mice (113.321857 ± 3.96 vs 56.27 ± 7.00 pg/mg, t=6.323, df=10, p<0.0001) (Fig. 3*a,b*). Conversely, the A313V show significantly elevated levels of homovanillic acid (HVA), increased by over 100% (42.99 ± 3.72 vs 136.94 ± 13.09 pg/mg, t=5.868, df=10, p=0.0002) (Fig. 3*c*), and increased ratios of DOPAC/DA (0.25 ± 0.037vs 0.56 ± 0.08 pg/mg, t=3.079, df=10, p=0.0117) and HVA/DA (0.09 ± 0.00 vs 1.20 ± 0.091 pg/mg, t=11.11, df=9, p<0.0001) (Fig. 3*d,e*). With the exception of DOPAC, these neurochemical measures replicate that which has been reported in DAT-KO mice, which completely lack DAT protein expression. While there is a 50% reduction of DOPAC levels in the mA313V mice, no changes in DOPAC levels were reported in DAT-KO mice (44).

**Figure 3.**
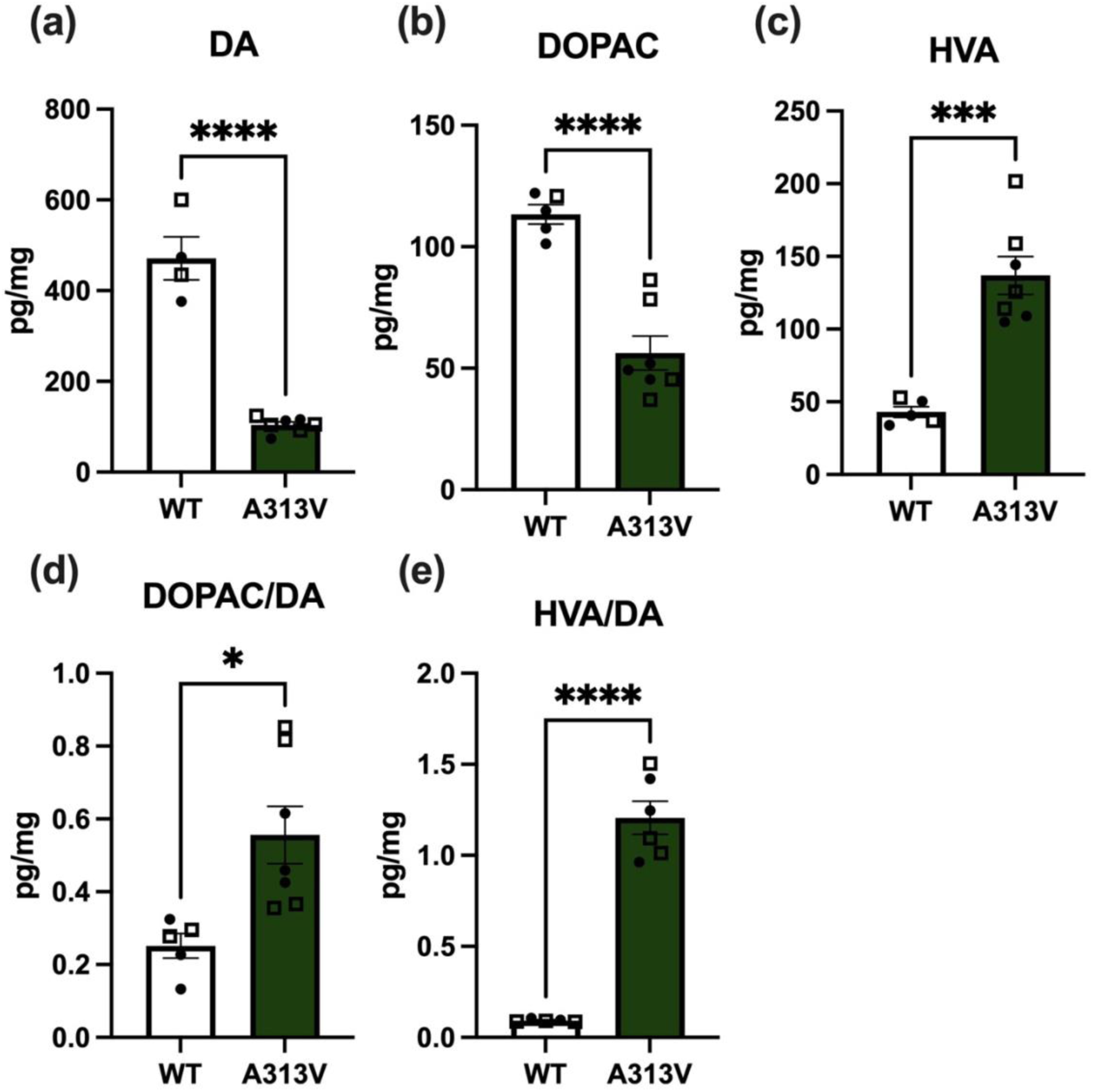
Striatal Tissue Content of Dopamine and Metabolites. HPLC analysis showed that levels of (a) dopamine (DA) (471.23 ± 47.51vs 104.13 ± 6.35 pg/mg, t=10.63, df=9, p<0.0001) and (b) DOPAC (113.321857 ± 3.96 vs 56.27 ± 7.00 pg/mg, t=6.323, df=10, p<0.0001) were greater, while (c) HVA (42.99 ± 3.72 vs 136.94 ± 13.09 pg/mg, t=5.868, df=10, p=0.0002) was lower in striatal homogenates from wildtype (WT) compared to dopamine transporter (DAT) A313V knock-in mice (A313V) (d) Ratios of DOPAC/DA (0.25 ± 0.037vs 0.56 ± 0.08 pg/mg, t=3.079, df=10, p=0.0117) and (e) HVA/DA (0.09 ± 0.00 vs 1.20 ± 0.091 pg/mg, t=11.11, df=9, p<0.0001) were lower in WT than A313V mice, indicating higher DA turnover in the A313V mice (n=4-7). All results are presented as mean ± SEM. Closed circular points represent values from female mice, and open square points represent values from male mice. Student’s unpaired, two-tailed *t-*tests, *p≤0.05, ***p≤0.001, ****p<0.0001

To test whether the reduced DA tissue content levels in the A313V mice were accompanied by a corresponding impairment in dopamine release and uptake, fast scan cyclic voltammetry was used to measure the rate of dopamine clearance from the extracellular space following electrically-evoked release in striatal slices (Fig. 4*a*). When DA clearance was fit to an exponential curve, A313V mice exhibited a longer decay time constant, tau, compared to WT mice (0.53 ± 0.02 vs 5.22 ± 0.17 seconds, t= 28.51, df=21, p<0.0001) (Fig. 4*b*). Indeed, the average tau value in A313V mice was approximately 10-fold greater than in WT mice. Moreover, the A313V mice have a near 80% reduction in V_max_ (4.24 ± 0.18 vs 0.76 ± 0.03 μM/second, t=18.61, df=21, p<0.0001) (Fig. 4*c*). Both phenotypes are indicative of decreased dopamine clearance, caused by lower levels of functional DAT as a result of the A313V variant.

**Figure 4.**
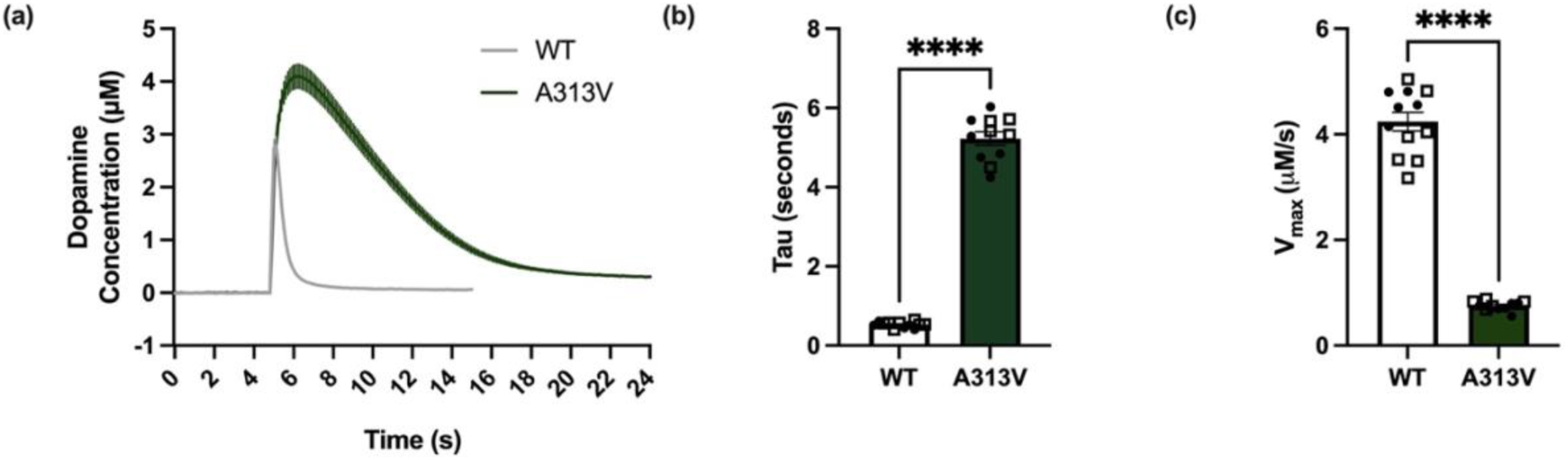
Dopamine kinetics measured in striatal slices from WT and A313V mice. Fast scan cyclic voltammetry was used to record dopamine responses to single electrical pulses (750 uA, 4 msec, biphasic) in dorsal striatal slices from wild-type (WT) and dopamine transporter (DAT) A313V knock-in mice (A313V). (a) Averaged raw dopamine concentration (μM) versus time in seconds (s) plots with SEMs from WT and A313V mice showed higher peaks and slower returns to baseline (clearance) in A313V mice (n=57-59). (b) Dopamine uptake as measured by the average time constant (tau), from an exponential decay model showed increased tau in A313V mice compared to WT mice (0.53 ± 0.02 vs 5.22 ± 0.17 seconds, t= 28.51, df=21, p<0.0001) (n=11-12). (c) The maximal velocity of dopamine uptake (Vmax), determined using the Michaelis-Menten model, was greater in WT compared to A313V mice (4.24 ± 0.18 vs 0.76 ± 0.03 μM/second, t=18.61, df=21, p<0.0001) (n=11-12). Closed circular points represent values from female mice, and open square points represent values from male mice. Bar graph results are presented as mean ± SEM. Student’s unpaired, two-tailed *t-*tests, ****p<0.0001

### Behavioral Characterization

Tight regulation of intra- and extra-cellular dopamine levels is critical for many cognitive and behavioral processes and can be affected by DAT levels. To examine the consequences of the A313V DAT variant on dopamine-dependent behaviors, a variety of behavioral tests were conducted. We first assessed body weight and observed no significant difference in weight between WT and A313V mice (25.08 ± 0.85vs 23.63 ± 0.70 grams, t=1.308, df= 39, p=0.1986) (Supp. Fig. 1a), unlike DAT-KO mice which are underweight as compared to WT animals (46).

The dorsal immobility response (DIR) — a stereotyped limb extension and immobility when mice are lifted by the nape of the neck — is shorter in duration when dopaminergic transmission is elevated (47). A313V mice tested with this assay were approximately three times as fast to move (decreased DIR) than their WT counterparts (64.89 ± 8.68 vs 26.80 ± 5.99 seconds, t=3.675, df=17, p=0.0019) (Fig. 5*i*), suggestive of a state of hyperdopaminergia due to reduced dopamine uptake. Consistent with this finding, when placed into a novel open-field arena for 150 minutes, adult A313V mice traveled over three times the mean total distance (total distance the subject has traveled; defines the subject’s location as the centroid of the subject) as their WT counterparts (3320.08 ± 153.95 vs 11303.43 ± 765.44 cm, t=9.873, df=52, p<0.0001) (Fig. 5*a,e*), and roughly two times greater mean horizontal activity (count of horizontal beam breaks) (15682.84 ± 519.67 vs 37977.06 ± 1561.93 beam breaks, t=12.72, df=52, p<0.0001) (Fig. 5*b,h*), vertical activity/rearing behavior (count of vertical beam breaks) (628.67± 44.97 vs 1238.28 ± 107.49 beam breaks, t=5.097, df=54, p<0.0001 (Fig.5*c,g*), and number of stereotypic behaviors (the number of beam breaks due to stereotypic activity; when the animal breaks the same beam or set of beams repeatedly) (9818.7 ± 493.73 vs 23804.04 ± 1079.89 beam breaks, t=11.63, df=53, p<0.0001) (Fig. 5*d,h*). When aged to 15 months, the A313V mice continued to be three-times as hyperactive compared to age-matched WT mice (3548.40 ± 648.59 vs 8659.40 ± 233.89 cm, t=7.413, df=8, p<0.0001) (Supp. Fig. 2).

**Figure 5.**
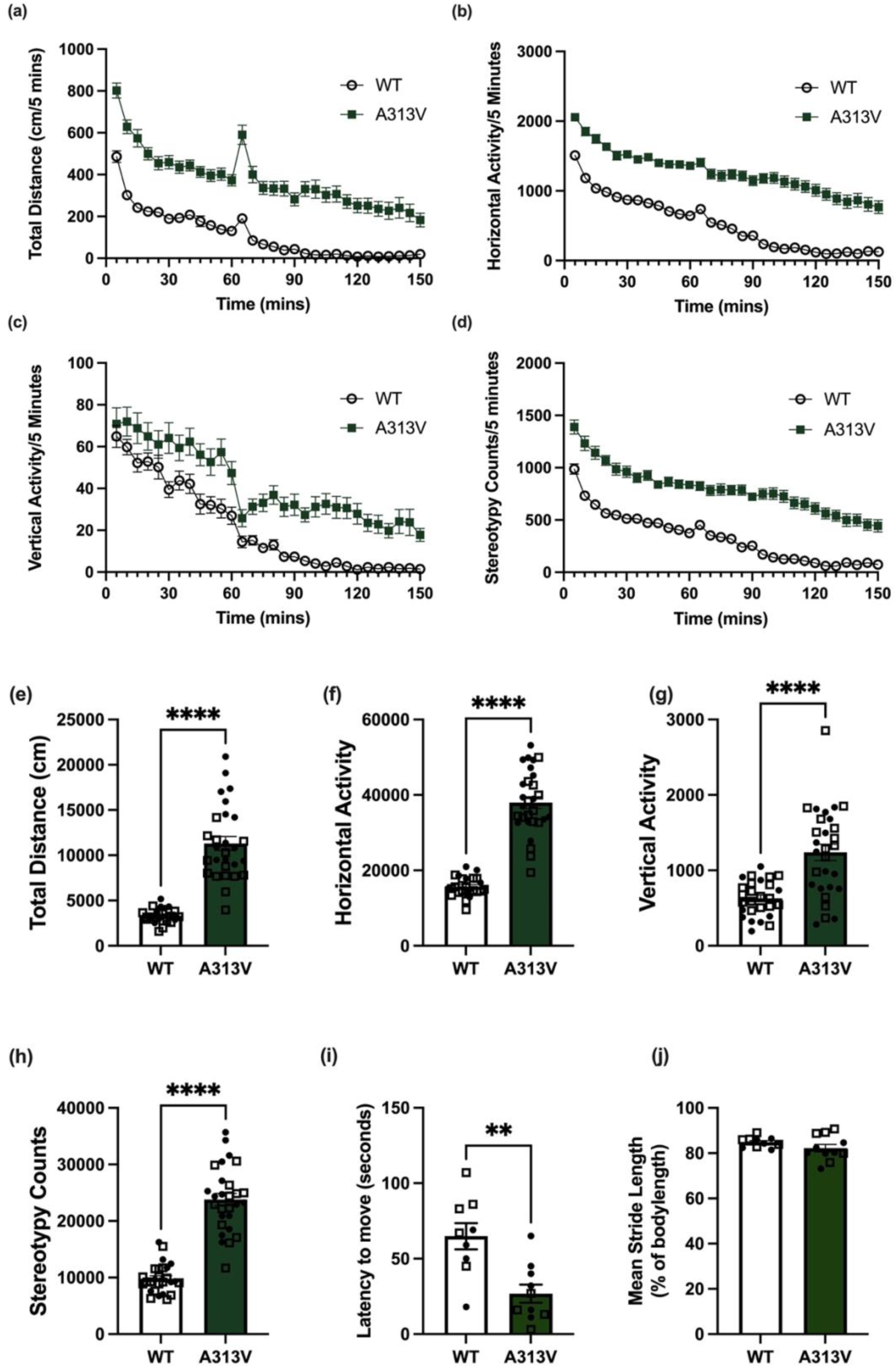
Baseline Locomotor Activity of adult WT and A313V mice. Baseline locomotor behavior of wildtype (WT), and dopamine transporter (DAT) A313V knock-in (A313V) mice. (a) Total distance, (b) vertical activity (c) horizontal activity, and (d) number of stereotypic counts were recorded in 5-minute bins for 150 minutes in an open field chamber. An i.p. injection of saline was given at 60 minutes. A313V mice showed significantly greater summed levels of (e) total distance (3320.08 ± 153.95 vs 11303.43 ± 765.44 cm, t=9.873, df=52, p<0.0001), (f) horizontal activity (15682.84 ± 519.67 vs 37977.06 ± 1561.93 beam breaks, t=12.72, df=52, p<0.0001), (g) vertical activity (628.67± 44.97 vs 1238.28 ± 107.49 beam breaks, t=5.097, df=54, p<0.0001), and (h) stereotypic count (9818.7 ± 493.73 vs 23804.04 ± 1079.89 beam breaks, t=11.63, df=53, p<0.0001) (n=25-29). (i) Latency to move in WT and DAT A313V mice was measured after clasping the animal by the nape of the neck to elicit the dorsal immobility response (DIR). A313V mice were significantly faster than WT mice to move (64.89 ± 8.68 vs 26.80 ± 5.99 seconds, t=3.675, df=17, p=0.0019) (n=9-10). (j) Mean stride length as a percentage of body length was assessed in WT and A313V mice. There was no significant difference in this measure between WT and A313V mice (84.84 +/- 0.73 vs 82.23 ± 1.68 mean stride length, t=1.369, df=19, p=0.1868) (n=10-11). Closed circular points represent values from female mice, and open square points represent values from male mice. Results are presented as mean ± SEM. Student’s unpaired, two-tailed t-tests were conducted, ns p>0.05, **p≤0.01, ****p<0.0001

Habituation of locomotor exploration was quantified using a habituation index (time in minutes to reach half of the maximal locomotor activity divided by total distance traveled using linear regression) (48). Habituation rate in A313V mice was roughly half that of WT controls (62.93 ± 0.72 vs 124.71 ± 13.75 minutes, t=4.402, df=51, p<0.0001) (Supp. Fig. 1b). This suggests that in addition to being hyperactive, the A313V mice have impaired locomotor habituation to a novel environment relative to WT mice.

A subset of DAT-KO mice, which fully lack DAT activity, have been reported to display gait abnormalities compared to WT mice, in the form of decreased stride length (49). We therefore performed gait analysis in A313V mice and mean stride length was measured as a percent of body length. Our results show that there is no difference in mean stride length between A313V and WT mice (84.84 +/- 0.73 vs 82.23 ± 1.68 mean stride length, t=1.369, df=19, p=0.1868) (Fig. 5*j*). Dopamine tone has also been shown to be modulatory in motor learning (50, 51), including in drosophila harboring the R445C DTDS-disease causing DAT variant, which exhibit impaired motor coordination (52). We next performed a nine-day rotarod experiment to measure both coordination (53) and motor learning (51). A mixed model analysis revealed no significant effect of genotype (F_1,13_=0.6625 p=0.4303), or a genotype by trial interaction (F_5,50_=1.727 p=0.1455). There was a significant effect of trial (F_5.65_ =14.79, p<0.0001) (Supp. Fig. 3a-c). In summary, the animals as a whole performed better over the consecutive trials. Rotarod performance was not significantly influenced by genotype or a genotype by trial interaction, suggesting no differences between A313V and WT animals throughout the experimental timeline.

### Symptom Management with Pharmacotherapy

While amphetamine (AMPH) injection induces hyperactivity in WT mice, AMPH decreases locomotor activity in DAT-KO mice (54). To address the hyperactivity symptoms observed at the early stages of DTDS, AMPH was administered to the A313V and WT mice and their locomotor activity was recorded. A two-way ANOVA was performed on the sum of total distance traveled post-treatment in WT and A313V mice. There were no main effects of genotype (F_1,18_=0.03197, p=0.8601) or treatment (F_1,18_=0.1693, p=0.6856), but there was a genotype by treatment interaction (F_1,18_=18.00, p=0.0005). Šidák’s multiple comparisons within genotype revealed a significant difference between WT mice treated with vehicle or AMPH (p=0.0112) and between A313V mice treated with vehicle or AMPH (p=0.0211). WT mice treated with 3 mg/kg AMPH showed an eightfold increase in locomotor activity compared to vehicle-treated WT animals (378.75 ± 64.65 vs 3190.20 ± 693.61 cm) (Fig. 6*a,c*). However, A313V mice treated with 3 mg/kg AMPH traveled approximately six times less than A313V mice that received vehicle (3698.60 ± 941.76 vs 568.86 ± 216.98 cm) (Fig. *6b,c*).

**Figure 6.**
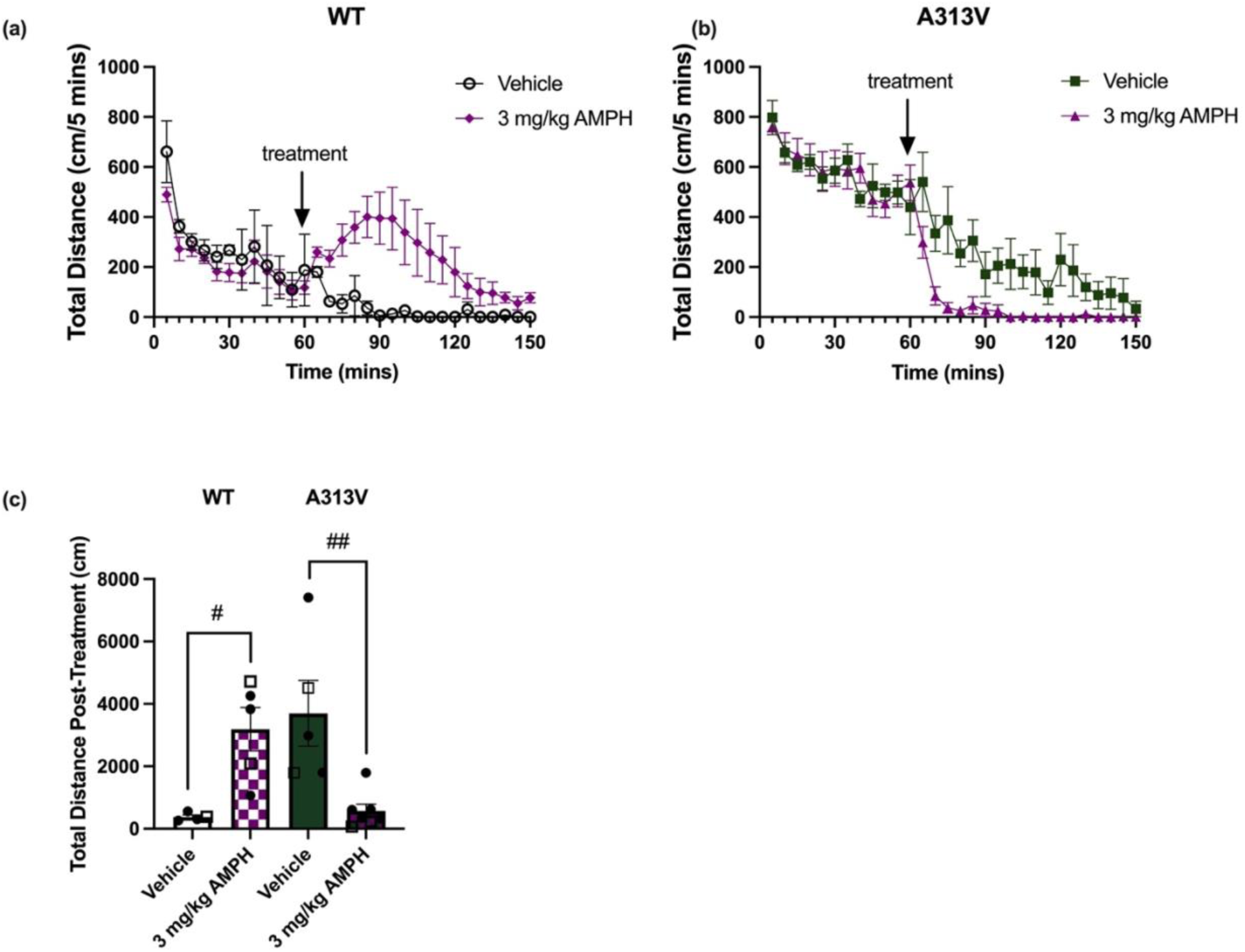
Amphetamine decreases locomotor activity in A313V mice. Total distance was recorded in 5-minute bins for 150 minutes in wildtype (WT) and dopamine transporter (DAT) A313V knock-in (A313V) mice. The test began with a 60-minute baseline followed by intraperitoneal (i.p.) administration of vehicle or amphetamine (AMPH) at the 60-minute timepoint. Drug intervention is marked with an arrow. (a) total distance traveled over time and (c) cumulatively in WT and (a,c) A313V mice treated with 3 mg/kg AMPH (n=4-7). A two-way ANOVA on the sum of total distance traveled post-treatment showed no main effects of genotype (F_1,18_=0.03197, p=0.8601) or treatment (F_1,18_=0.1693, p=0.6856). There was a genotype by treatment interaction (F_1,18_=18.00, p=0.0005). WT animals treated with AMPH traveled more than those treated with vehicle (378.75 ± 64.65 vs 3190.20 ± 693.61 cm, p=0.0112), whereas A313V animals treated with AMPH traveled significantly less than those treated with vehicle (3698.60 ± 941.76 vs 568.86 ± 216.98 cm, p p=0.0211), as revealed by Šidák’s multiple comparisons within genotype. In bar graphs, closed circular points represent values from female mice, and open square points represent values from male mice. Results are presented as mean ± SEM. A two-way ANOVA examining treatment, genotype, and interactions, with Šidák’s multiple comparisons was performed. For post hoc effects: #p<0.05, ##p≤0.01

Next, we assessed the effects of methylphenidate (MPH), which has also been shown to reduce hyperactivity of DAT-KO mice similarly to AMPH (54). Moreover, MPH and AMPH are both clinically used for the treatment of ADHD. Interestingly, MPH administration was not able to decrease hyperactivity in the A313V mice compared to vehicle-treated animals, as revealed by a two-way ANOVA with Tukey’s multiple comparisons performed on the sum of total distance traveled post-treatment. There was a significant main effect of treatment (F_2,79_=30.25, p<0.0001), genotype (F_1,79_=17.31, p<0.0001), and a significant genotype x treatment interaction (F_2,79_=10.59, p<0.0001). Tukey’s multiple comparisons showed that, in WT mice, total distance traveled upon treatment of both 30 mg/kg and 40 mg/kg was increased compared to vehicle (652.62 ± 63.32 vs 8156.67 ± 750.17 cm, p<0.0001; 652.62 ± 63.32 vs 4468.40 ± 1127.82, p=0.0020). For A313V mice, there was no significant differences between vehicle and 30 mg/kg MPH (5658.48 ± 576.75 vs 7357.29 ± 535.62 cm, p = 0.1693), and vehicle and 40 mg/kg MPH (5658.48 ± 576.75 vs 7623.44 ± 715.31 cm, p = 0.0581) (Supp. Fig. 4a-h). This observation is in contrast to what has been reported in DAT-KO mice: treatment with 30 mg/kg of MPH decreased locomotor activity in DAT-KO mice compared with vehicle controls (54).

Tyrosine hydroxylase (TH) is the rate limiting enzyme in the synthesis of DA, and inhibition of TH with alpha-methyl-para-tyrosine (ɑMPT) greatly reduces the behavioral hyperactivity of DAT-KO mice (55). Therefore, the effects of ɑMPT on A313V and WT mice in the open-field arena were assessed at doses of 250 mg/kg, 125 mg/kg, 62.5 mg/kg, and 31 mg/kg. A two-way ANOVA conducted on the sum of total distance post-treatment showed a main effect of treatment (F_3,74_=6.690, p=0.0005) and genotype (F_1,74_=9.174, p=0.0034), and a genotype x treatment interaction (F_3,74_=7.401, p=0.0002). Multiple comparisons within genotype showed that compared to vehicle, neither of the four doses of ɑMPT lead to significant changes in the sum total distance post-treatment in WT mice (Fig. 7*a,c,e*) (652.62 ± 63.32 vs 559.00 ± 172.21 cm, p=0.994; 652.62 ± 63.32 vs 1320.67 ± 222.12 cm, p=0.9418; 652.62 ± 63.32 vs 986.33 ± 194.15 cm, p=0.9920). In contrast, A313V mice treated with 250 mg/kg ɑMPT traveled five-fold less than those treated with vehicle (5658.48 ± 576.75 vs 1032.29 ± 215.05 cm, p<0.0001). Moreover, both doses of 125 mg/kg (5658.48 ± 576.75 vs 2240.00 ± 44.60, p=0.0239) and 62.5 mg/kg (5658.48 ± 576.75 vs 2171.00 ± 186.81, p=0.0204) also decreased total distance traveled by over half (Fig. 7*b,d*). Treatment with 31 mg/kg ɑMPT led to a main effect of genotype (F_1,19_=43.81, p<0.0002), no main effect of treatment (F_1,19_=3.241, p=0.0877), and a significant genotype by treatment interaction (F_1,19_=6.445, p=0.0200). Šidák’s multiple comparison test showed that compared to vehicle, the sum of total distance post-treatment was decreased in A313V mice that received 31mg/kg ɑMPT (5197.34 ± 912.52 vs 3033.25 ± 503.64 cm, p=0.0147), but there was no difference between treatments in WT animals (629.97 ± 122.53 vs 998.15 ± 276.48 cm, p=0.8391) (Fig. 7*e*).

**Figure 7.**
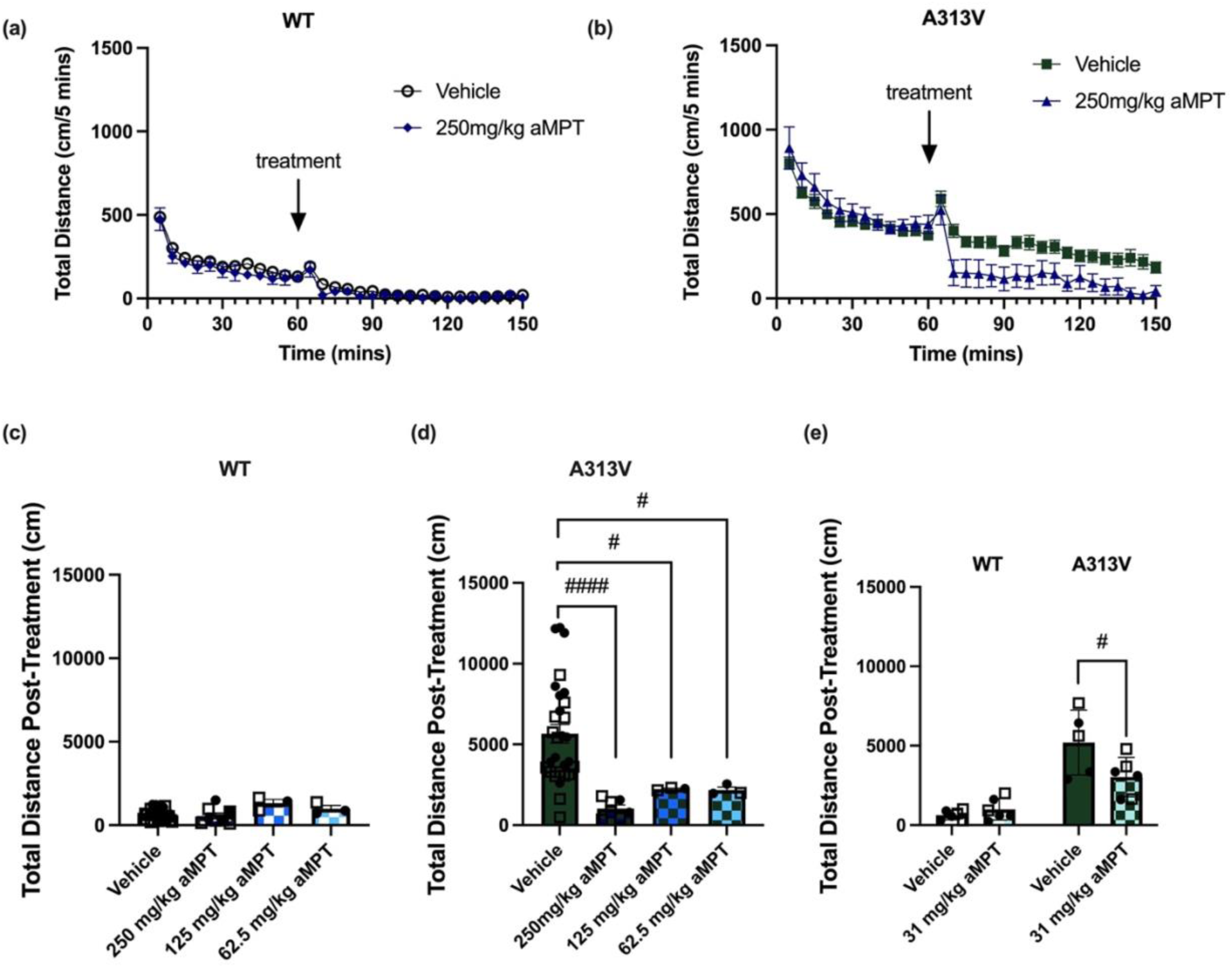
Analysis of Four Different Doses of Alpha-methyl-para-tyrosine in WT and A313V Mice. Total distance was recorded in 5-minute bins for 150 minutes in wildtype (WT) and dopamine transporter (DAT) A313V knock-in (A313V) mice. The test began with a 60 minute baseline followed by intraperitoneal (i.p.) administration of vehicle or alpha-methyl-para-tyrosine (ɑMPT) at the 60-minute timepoint. (a) Total distance over time in WT and (b) A313V mice treated with vehicle or 250mg/kg ɑMPT (n=8-28). An arrow indicates the time point at which treatment was administered. This graph is representative and illustrates the temporal pattern of ɑMPT administration and effect for one dose. (c,d) Bar graph displaying the effects of vehicle or decreasing doses of ɑMPT at 250 mg/kg, 125 mg/kg, 62.5 mg/kg, and (e) 31 mg/kg on the sum of total distance traveled post-treatment. A two way-ANOVA with Tukey’s multiple comparisons showed that A313V mice treated with 250 mg/kg (5658.48 ± 576.75 vs 1032.29 ± 215.05 cm, p<0.0001), 125 mg/kg (5658.48 ± 576.75 vs 2240.00 ± 44.60, p=0.0239), and 62.5 mg/kg (5658.48 ± 576.75 vs 2171.00 ± 186.81, p=0.0204) ɑMPT traveled less than those treated with vehicle, whereas there were no such changes seen in WT mice. A separate two-way ANOVA with Šidák’s multiple comparisons showed similar decreases in total distance traveled in A313V mice treated with 31 mg/kg ɑMPT (5197.34 ± 912.52 vs 3033.25 ± 503.64 cm, p=0.0147), but not in WT mice. Closed circular points represent values from female mice, and open square points represent values from male mice. Results are presented as mean ± SEM. For post hoc effects: #p≤0.05, ###p≤0.001.

### Disease modification using pharmacological chaperones

It has been shown that pharmacological chaperones of DAT, including ibogaine and noribogaine, can rescue expression of select DTDS disease causing variants in cells (30, 31, 41). Moreover, noribogaine was shown to rescue motor and sleep impairments of the G108Q, V158F, and G327R DTDS variants in drosophila (31, 41). In order to assess whether pharmacological chaperones could also rescue the A313V variant *in vivo* in a mammalian system, a series of experiments were performed with noribogaine, one of the most efficacious DAT pharmacological chaperones. Cell based studies have shown that high doses of noribogaine (50-100μM) are needed to achieve pharmacological chaperone activity (30). Thus, pharmacokinetic (PK) studies were first conducted to establish the proper dosing regimen that would allow achievement of the 50-100μM brain concentration needed for pharmacological chaperone activity. Detailed PK studies were conducted via oral gavage based on experiments conducted by (56), as ICV injections of noribogaine could not achieve brain doses in the range needed for our studies (data not shown). As a first step, mean noribogaine concentrations were measured by LC/MS/MS in cerebellar brain tissues at 2 and 24 hours after oral administration of 100 mg/kg. As shown in Supp. Fig 5a, this dose resulted in 27.94 μM and 2.06 μM of noribogaine in the brain after 2 and 24 hours respectively; which are below the concentrations needed for pharmacological chaperone activity (Supp. Fig. 5a). The dosing regimen was therefore modified, and another cohort of mice received 500 mg/kg of noribogaine via oral gavage followed by a second 250 mg/kg dose (oral gavage) 5 hours later. As shown in Supp. Fig 5b, noribogaine levels were above 100 μM (∼180-200 μM) at 1 hour and following the second injection remained above this for at least 24 hours (Supp. Fig. 5b). Thus, a dosing regimen of 500 mg/kg noribogaine via oral gavage followed by a second dose of 250 mg/kg five hours later is sufficient to achieve 100 μM noribogaine in the brain that is largely stable across at least 19 hours.

Next, it was important to determine the effective duration that noribogaine concentrations in the brain remained above 100 μM. The same dosing regimen (500 mg/kg noribogaine via oral gavage followed by 250 mg/kg five hours later) was carried out and plasma was collected at different time points (1, 5, 6, 12, 24, 48, 72, 80, 96, 120, and 144 hours) after the initial 500 mg/kg noribogaine dose. Brains were also collected at the 72, 80, 96, 120, 144, and 168 hour timepoints. After the 500+250 mg/kg oral gavage, brain noribogaine concentrations fell below 100 μM at approximately 24 hours and below 50 μM at approximately 36 hours after first oral gavage (Supp. Fig. 5c). Regression analysis of plasma vs. brain μM noribogaine from the experiment in Supp. Fig 5b and Supp. Fig. 5c gave an R-squared value of 0.8731, suggesting that plasma noribogaine concentration can be used to estimate noribogaine concentrations in the brain (Supp. Fig. 5d).

A fourth experiment explored the effects of repeated noribogaine dosing on plasma and brain noribogaine concentrations. Here, the above 500 + 250 mg/kg noribogaine schedule was administered every 48 hours over the course of 9 days, with brains collected on day 10, 24 hours after the 4th noribogaine dose. Blood was also collected from two groups of animals on day 2 (group 1) and 4 (group 2), 24 hours after the 500 mg/kg noribogaine administration on days 1 and 3, respectively. The experiment timeline is illustrated in Supp. Fig. 6a. The mean noribogaine plasma concentration on day 2 was 30.93 μM and on day 4 was 21.23 μM (Supp. Fig. 6b). The mean noribogaine brain concentration on day 10 was 35.68 μM (Supp. Fig. 6b). The brain concentrations of noribogaine were estimated based on the previously established relationship between plasma and brain noribogaine concentrations. This indicated that on day 2, brain noribogaine concentrations were approximately 169.52 μM and 116.21 μM on day 4. This experiment suggests that following repeated dosing 48 hours apart, noribogaine levels in the brain fall below 100 μM at approximately 96 hours (4 days, Supp. Fig. 6c).

Having optimized the noribogaine administration paradigm to achieve 100μM concentrations in the brain, an experiment was conducted to assess whether the ER-retained A313V DTDS variant could be rescued with noribogaine. The experiment was carried out over the course of four days (Fig. 8*a*). The noribogaine dosing combination of 500 mg/kg + 250 mg/kg was administered twice, 48 hours apart, on day 1 and day 3. Locomotor activity was assessed 24 hours after noribogaine administration on day 4, and brains were collected upon completion of the open field test. A two-way ANOVA was conducted on the sum of total distance traveled. There was a main effect of treatment (F_1,25_=22.95, p<0.0001) and genotype (F_1,25_=39.24, p<0.0001), but no interaction effect (F_1,25_=2.356, p=0.1374), suggesting that treatment with noribogaine decreased levels of locomotor activity similarly in both WT (3687.86 ± 336.01 vs 1912.71 ± 311.13 cm) and A313V (7939.86 ± 836.61 vs 4491.25 ± 525.87) mice (Fig. 8*b-e*). Western blots were then performed on striatal and midbrain tissue to assess levels of DAT in animals treated with vehicle or noribogaine. In WT animals, the difference in levels of DAT in the striatum (100.00±35.81 vs 65.82 ± 10.09 % of WT vehicle-treated average, t=0.8542, df=11, p=0.4112) and midbrain (100.00 ± 14.67 ± 70.96 ± 12.08 % of WT vehicle-treated average t=1.544, df=9, p=0.1570) were not significantly different between vehicle and noribogaine treated animals (Fig. 9*a-d*). Similarly, there was no difference in the levels of DAT in the striatum (100.00 ± 15.90 vs 88.65 ± 6.17 % of A313V vehicle-treated average, t=0.7063, df=11, p=0.4947) or midbrain (100.00 ± 61.88 vs 51.74 ± 17.28 % of A313V vehicle-treated average,t=0.7511, df=10, p=0.4699) in A313V mice treated with vehicle or noribogaine (Fig. 9*e-h*). This result shows that, although this noribogaine regimen can reduce hyperactivity of both WT and A313V animals, it does not increase DAT expression, which is in contrast to what has been reported in cell-based studies.

**Figure 8.**
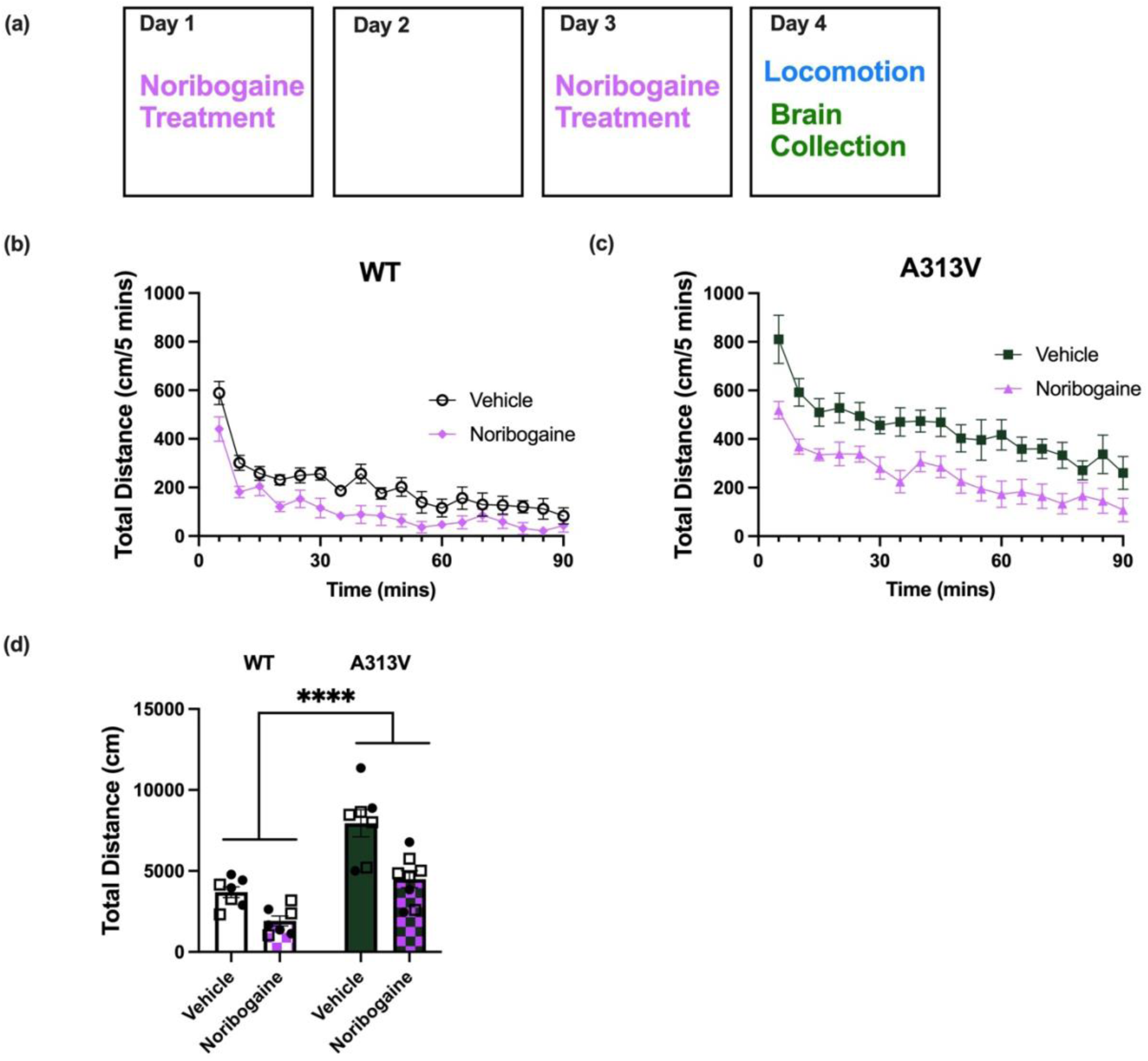
Chronic Noribogaine Administration Reduces Locomotor Activity in WT and A313V mice. (a) Noribogaine was administered via oral gavage according to the established dosing regimen of 500 mg/kg followed by 250 mg/kg five hours later (“Noribogaine Treatment”). This dosing regimen was done on day 1 and on day 3, 48 hours apart. Locomotor activity was assessed 24 hours after the final dose (day 4, Locomotion). (b) Total distance traveled over time and cumulative distance in wildtype (c) (WT) and (c, d) A313V mice treated with vehicle or noribogaine. A two-way ANOVA found a main effect of treatment (F_1,25_=22.95, p<0.0001) and genotype (F_1,25_=39.24, p<0.0001), but no interaction effect (F_1,25_=2.356, p=0.1374), suggesting that treatment with noribogaine decreased levels of locomotor activity similarly in both WT (3687.86 ± 336.01 vs 1912.71 ± 311.13 cm) and A313V (7939.86 ± 836.61 vs 4491.25 ± 525.87) mice. Closed circular points represent values from female mice, and open square points represent values from male mice. Bar graphs results are presented as mean ± SEM. A two-way ANOVA with Šidák’s multiple comparisons was performed. For main effects: ****p<0.0001

**Figure 9.**
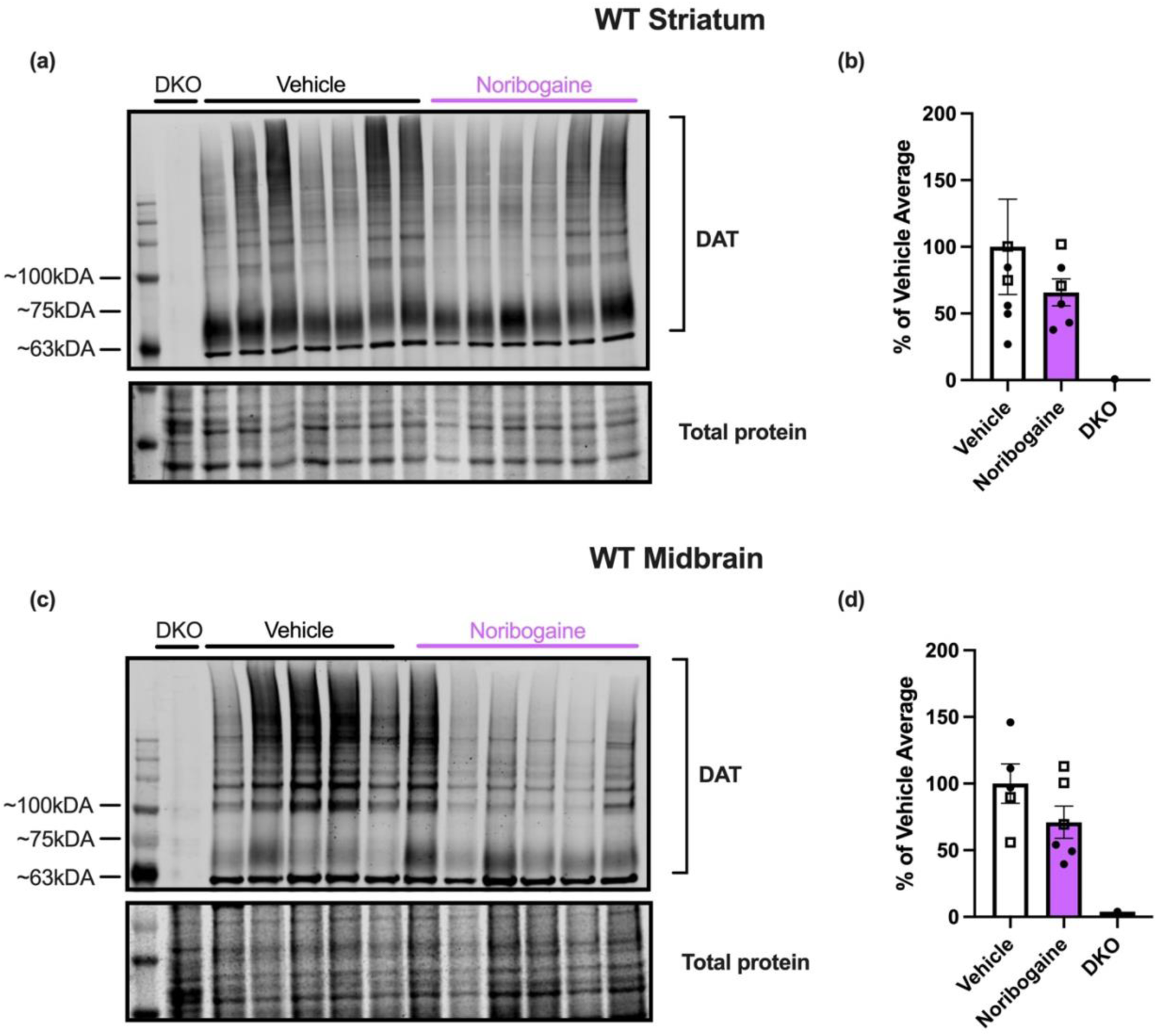
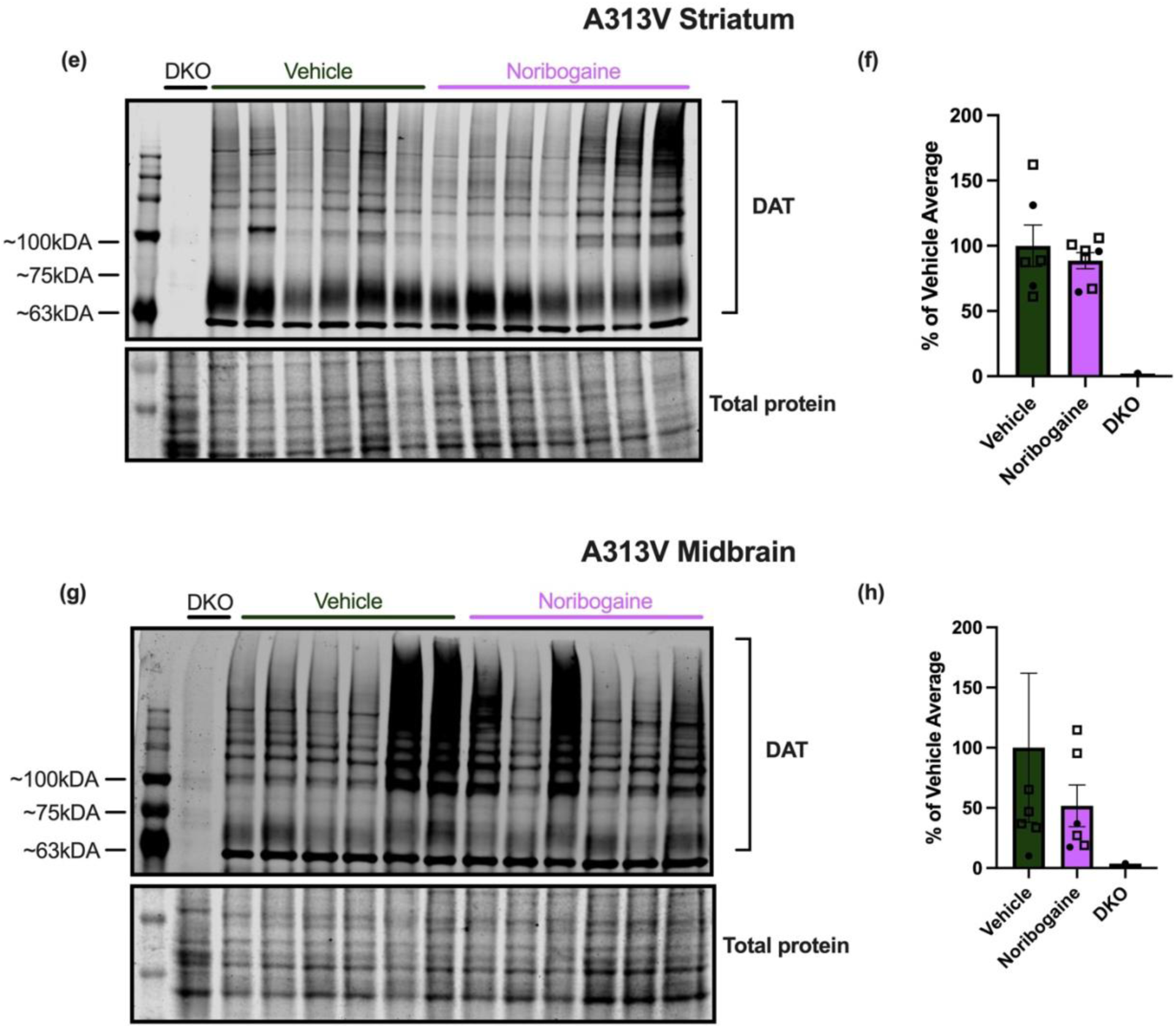
DAT expression levels in the striatum and midbrain of WT and A313V mice after treatment of noribogaine or vehicle. Dopamine transporter (DAT) levels were assessed in wild-type (WT) and A313V DAT knock-in (A313V) mice in the (a-b, e-f) striata and (c-d, g-h) midbrains of animals treated with vehicle or noribogaine. DAT-knockout (DKO) mouse samples were used as a negative control to confirm antibody specificity and to identify the appropriate regions for DAT analysis. The lower panels show a cropped total protein stain, used here to visualize approximately even protein loading. Lanes are labeled to indicate DKO, vehicle-treated, and noribogaine-treated samples. Molecular weight is shown in kDA to the left. DAT signal intensity was normalized to the total protein loading control and is expressed as a percentage of the mean vehicle-treated value. There was no statistically significant difference in DAT levels in the striatum of WT (100.00±35.81 vs 65.82 ± 10.09 % of WT vehicle-treated average, t=0.8542, df=11, p=0.4112) or A313V (100.00 ± 15.90 vs 88.65 ± 6.17 % of A313V vehicle-treated average, t=0.7063, df=11, p=0.4947) mice between animals treated with vehicle or noribogaine. There was no statistically significant difference in DAT levels in the midbrain of WT (100.00 ± 14.67 ± 70.96 ± 12.08 % of WT vehicle-treated average, t=1.544, df=9, p=0.1570) or A313V (100.00 ± 61.88 vs 51.74 ± 17.28 % of A313V vehicle-treated average, t=0.7511, df=10, p=0.4699) mice between animals treated with vehicle or noribogaine. Closed circular points represent values from female mice, and open square points represent values from male mice. Bar graphs results are presented as mean ± SEM. Student’s unpaired, two-tailed *t-*test was conducted separately for WT and A313V mice. n.s. p>0.05

## Discussion

Because folding efficiency is not 100% even for most wild-type proteins (57), a single amino acid change is often sufficient to induce disease pathogenesis by leading to protein misfolding (58). Indeed, misfolding of membrane proteins is linked to a number of diseases such as nephrogenic diabetes insipidus, cystic fibrosis, Fabry disease, and hypogonadotropic hypogonadism. Misfolding of eukaryotic membrane proteins typically occurs in the endoplasmic reticulum, where the misfolded protein is retained and targeted for degradation (28).

Dopamine transporter deficiency syndrome (DTDS) is believed to be caused by an analogous mechanism wherein variations in SLC6A3 lead to DAT protein misfolding, retention in the ER, and consequent low levels of functional DAT in striatal nerve terminals. Indeed, DAT is synthesized in the cell bodies of dopamine neurons within the midbrain and then trafficked to nerve terminals in the striatum (59, 60). In the initial part of this study, the neurochemical phenotype of the DAT A313V DTDS mice was examined, which models the human A314V DTDS variant. Firstly, results showed that the A313V DAT variant indeed leads to the retention of the immature, core glycosylated DAT protein in the midbrain, coincident with a decrease in mature, fully glycosylated DAT in the striatum. These show that the mechanism by which the A313V DAT variant leads to DTDS is due to 1.) its misfolding and retention in the ER, and 2.) the consequent low levels of functional DAT needed to maintain normal, physiological function.

Levels of dopamine metabolites, including HVA, are increased in cerebral spinal fluid of human patients with DTDS (15–17) and these changes in dopamine metabolites are one of the diagnostic criteria for the disease (27). The A313V knock-in mice recapitulate this neurochemical change, as observations also demonstrate increased amounts of HVA in these animals compared to WT mice. Furthermore, these results show that there is decreased tissue levels of DA in the A313V mice, consistent with the hypothesis that low levels of functional DAT led to decreased DA recycling and ultimately reduced DA tissue levels in the A313V mice. Such reduction in DA tissue levels is also observed in DAT-KO mice (44, 45). Interestingly, results show that the A313V mice had lower levels of DOPAC compared to WT mice yet increased levels of HVA. This is consistent with DOPAC representing mostly intracellular dopamine metabolism by MAO, (61, 62), while HVA primarily reflects metabolism by *extraneuronal* catechol-O-methyltransferase (COMT) in glial cells (62–64) potentially after transport into the cell by the organic cation transporters (65).

The principal role of the DAT is to spatially and temporally modulate DA neurotransmission through uptake of DA from the extracellular space after release, where it can then be repackaged into vesicles to be released again. Low levels of DAT lead to decreased DA uptake, consequently increased DA levels in the extracellular space, and thus increased dopaminergic neurotransmission. The reported initial behavioral symptoms of patients with DTDS are reflective of this hyperdopaminergia, characterized by features of chorea, ballismus, and orolingual dyskinesia (27). The behavioral characterization of the A313V mice recapitulates this hyperkinetic stage of the disease. Results showed that the A313V mice have decreased dorsal immobility response (DIR) time, and are hyperactive in the open field test compared to their WT littermates. These are both proxies for increased dopaminergic neurotransmission in mice.

Importantly, results suggest that two compounds are able to reduce the hyperactivity seen in the A313V mice: amphetamine (AMPH) and alpha-methyl-para-tyrosine (ɑMPT). Both of these drugs are currently approved for use in human patients and could be of translational interest for DTDS patients (66, 67). Of particular interest is ɑMPT because of its potential to be a disease-modifying treatment for DTDS. The mechanism of action of ɑMPT is inhibition of conversion of tyrosine to L-DOPA by tyrosine hydroxylase, therefore blocking the production of dopamine (67). This effectively lowers the amount of dopamine neurotransmission, which in turn reduces locomotor activity of the A313V mice. It is well known that both intracellular and extracellular dopamine can induce toxicity (49, 68–73) and up to 40% of dopamine transporter knock-out (DAT-KO) mice develop gait abnormalities and tremor due to loss of GABAergic striatal medium spiny neurons (49). Importantly, in DAT-KO mice, chronic treatment with ɑMPT was able to ameliorate the behavioral manifestation of this neurodegeneration and reduce mortality rate over a 40-week period to 0% (49). Considering the similar effects we observed on the motor activity of A313V mice after ɑMPT treatment, we believe that ɑMPT has the potential to modify the disease course of DTDS and reduce disease burden when used during the initial stages of the disease. As mentioned previously, ɑMPT is currently used in humans to treat symptoms associated with pheochromocytoma at doses of 600-3500 mg daily (67). Side effects at these doses include sedation, diarrhea, anxiety, nightmares, crystalluria, galactorrhea, and extrapyramidal symptoms (74). However, Ankenman & Salvatore, 2007 suggest that at low doses these side effects can be mitigated. We show in our studies that ɑMPT can dose-dependently reduce hyperactivity of A313V mice with doses as low as 31 mg/kg. This suggests that low to moderate dosing of ɑMPT could also have potential clinical benefit for DTDS patients with potentially modest adverse effects.

Ibogaine and noribogaine have been shown to act as pharmacological chaperones of DAT and rescue the A314V DTDS disease causing variant in cells (30, 31, 41, 43). With this in mind, experiments were conducted with noribogaine and assessed its potential efficacy as a disease modifying treatment to rescue the levels of the ER-retained A313V DAT. Although both WT and A313V animals treated with noribogaine showed decreased levels of locomotor activity, no changes were observed in DAT protein levels as assessed by western blots in the striatum or midbrain in WT and A313V animals treated with vehicle or noribogaine. This result indicated that under the current experimental conditions, it was not possible to observe rescue of A313V DAT protein expression i*n vivo* with noribogaine. There could be several reasons explaining the lack of effect of noribogaine. The simplest explanation could be that noribogaine is not able to act as a pharmacological chaperone of DAT *in vivo* in a mammalian model, in contrast to what has previously been described in cell-based systems. Alternatively, it is possible that the analysis of DAT levels with western blots is not sensitive enough to detect minor upregulation of DAT protein. Lastly, it is possible that the right dosing regimen of noribogaine was not achieved to observe its pharmacological chaperoning effect *in vivo*, in mice. This may, however, be unlikely, as extensive PK studies were performed to ensure that proper levels of noribogaine were attained the brain. What remains puzzling is that (31) demonstrated the ability to detect expression of human DAT G140Q or V158F variants in axonal compartments of drosophila when flies were fed with noribogaine beginning in the instar larval stage throughout adulthood. Interestingly, (41) showed that feeding adult drosophila with noribogaine for only 5 days led to “substantially” lower levels of hDAT G140Q variant in axonal compartments. It is therefore possible that, in our studies, where noribogaine was administered to the A313V mice only in adulthood, rescue was not achieved and perhaps treatment with noribogaine needs to begin earlier in life, and for a longer duration, to observe DAT rescue. However, this is difficult to experimentally verify because noribogaine levels in the brain fall below the efficacious 100 μM levels after approximately 84 hours of administration, despite repeated dosing every 48 hours. Therefore, it may not be pharmacokinetically feasible to conduct prolonged noribogaine studies with early intervention in mice.

As mentioned previously, the second stage of DTDS is characterized by Parkinson’s-like symptoms, manifesting as dystonic posturing, resting and action tremor, bradykinesia, and rigidity in humans. In the A313V animals, we did not observe or report any “Parkinson’s disease-like” phenotypes, even when motor activity was assessed in aged 15-month old animals. The only motor symptom that could be consistently observed was that of hyperkinesia, which mimics the early stage of DTDS. One potential reason for this may be that mice have a substantially greater capacity to cope with DA depletion than humans, otherwise known as a greater motor reserve. Indeed, developmental ablation of nearly 90% of ventral tegmental area and substantia pars compacta dopaminergic neurons in mice produces no notable motor phenotype (75), while it is known that in humans, the clinical motor symptoms of Parkinson’s Disease emerge when approximately 50-60% of substantia nigra axon terminals have been lost (76). Therefore, it may not be possible to capture the dystonic late-stage parkinsonian symptoms of DTDS in a mouse model, limiting insights solely to the initial stage of the disease.

Alternatively, the parkinsonism symptoms of DTDS may arise through an orthogonal mechanism related to ER retention of DAT. Patients harboring other SLC6A3/DAT variants that are not hypothesized to be ER-retained such as T356M (11), A559V (12, 77), and E602G (77) do not show symptoms of parkinsonism. Further, young-onset Parkinson’s disease is treated with levodopa/dopamine therapy (78,79), while DTDS patients do not respond to this treatment (15–17); this suggests a potential alternative pathology by which DTDS parkinsonism-dystonia arises. However, neuropathology analyses have not yet been performed in DTDS patients to elucidate if there is neurodegeneration in these patients, and, if so, in which neuronal population(s).

In summary, we have recapitulated some of the key DTDS symptoms and neurochemical changes seen in humans in our A313V-DAT mouse model. Furthermore, we have provided evidence for two treatments for the hyperkinetic symptoms of the disease: AMPH and ɑMPT. Importantly, we believe that ɑMPT may serve as a disease-modifying treatment for DTDS by directly addressing the increased dopaminergic tone seen in the disease. Overall, this study presents a new model of DAT hypofunction that displays a unique set of phenotypes distinct from DAT-KO mice, identifying potential translational biomarkers relevant to dopamine-related disorders, such as DTDS.

## Materials and Methods

### Mice and genotyping

Experiments were conducted in adult mice. Mice within a genotype were arbitrarily assigned to an experimental group so that each group contained approximately equal numbers of males and females. Sex was not treated as a biological variable and data from male and female mice were combined. Mice were housed with littermates on a 12-hour light-dark cycle with food and water available *ad libitum*. All experimentation was conducted during the light phase. *SLC6A3*^A313V+/+^ (A313V) mice were created via CRISPR-Cas9 endonuclease-mediated transgenesis. Targeting was performed on an FVB/N × C57BL/6J hybrid F1 background using an sgRNA designed to a 20bp target within Exon 7 of *Slc6a3* (target sequence: GGAGAAGCACACCTGGGTGG). A 127bp ssODN was provided for homology-directed repair to introduce the desired nucleotide change (NM_010020.3:c.1095C>T; p.Ala313Val) along with a second silent nucleotide change made to facilitate targeting (NM_010020.3:c.1093T>C ; p.Asp312Asp) (ssODN sequence: (CGGGTGGAACTAACTCACATCCCATTCTCTCTGATCATCTTCCTCATGCTCTCCCATCTCTGGCTGTGACAGGTGTGGATCGACGTCGCCACCCAGGTGTGCTTCTCCCTTGGCGTTGGGTTTGGGG). Founder mice were identified by Sanger sequencing and bred to a C57BL/6J background to generate wildtype (WT) and A313V mice. Following germline transmission, A313V mice were backcrossed onto the C57Bl/6J background, and continuously backcrossed for at least 9 generations prior to starting experimentation. Mice were genotyped using two separate PCRs for each allele: the WT and A313V variant allele amplification primers (552-bp amplicon) (along with a control reaction for PCR); WT forward: ACA GGT GTG GAT CGA TGC, reverse: GGA GAG AGG ATG GAG AAG AGA A; A313V forward: ACA GGT GTG GAT CGA CGT, reverse: GGA GAG AGG ATG GAG AAG AGA A. Both reactions are run side-by-side. Touchdown PCR was used with DreamTaq DNA Polymerase (Thermo Fisher Scientific): 2 min at 96 degrees, then each cycle after is 94 degrees for 20 seconds, annealing for 30 seconds. The initial 12 cycles are touchdown from 64-58 degrees (0.5 degrees/cycle), with the remaining 28 cycles with an annealing temperature of 58 degrees (40 cycles total), with extension of 30 seconds for each cycle. Animals were genotyped prior to experiments and after experiments were completed to confirm the correct genotype.

### Drugs

**Table 1.** Drugs.

| Drug | Source |
| --- | --- |
| Amphetamine | Toronto Research Chemicals, #A634295 |
| Threo-Methylphenidate hydrochloride (MPH) | Tocris Bioscience, #298-59-9 |
| alpha-methyl-D,L-p-tyrosine, Methyl Ester (αMPT) | ChemCruz, Santa Cruz Biotechnology, Inc. #sc-219470 |
| Ibogaine | Ibogaworld, ibogaworld.com |
| Noribogaine | Synthesized from ibogaine by Dalriada Drug Discovery, Mississauga, ON, CA, and by Dr. Landon Edgar at the University of Toronto, Toronto, ON, CA. |

#### Noribogaine HCl Synthesis

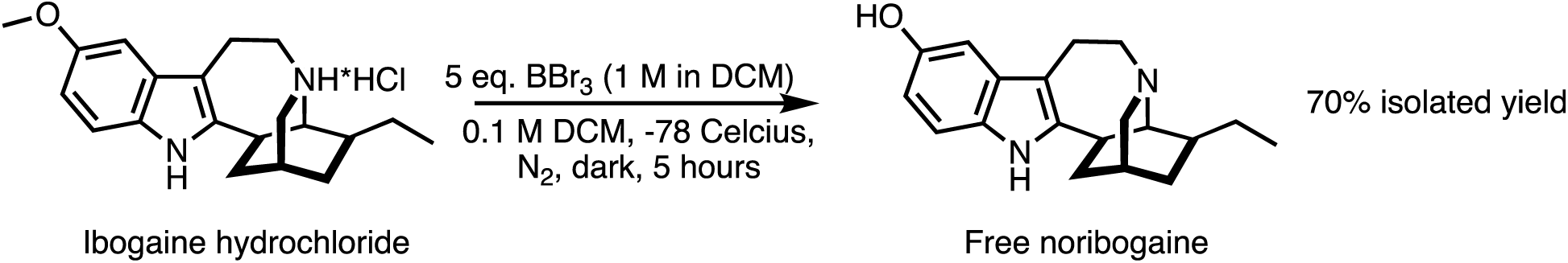

Ibogaine hydrochloride (3g, 9.66mmol) was dissolved in 100mL of anhydrous DCM (0.1 M). The reaction flask was purged with nitrogen and cooled to -78^°^C before 1M solution of boron tribromide in DCM (5 eq., 48.32mmol) was added dropwise over 20 minutes. The reaction was left stirring in the dark for 5 hours. The crude was poured slowly into sat. aq. NaHCO_3_ and extracted 200 mL of 15% MeOH in CHCl_3_ three times. The aqueous layer was diluted with aq. NH_4_OH and extracted successively with 200 mL EtOAc four times. The organic fractions were combined, dried over MgSO_4_, filtered, and concentrated *in vacuo*. The organic fractions were then purified by silica gel flash column chromatography (3%-15% MeOH in DCM) to yield pure noribogaine free base (4.08 g, 70% yield). **TLC:** (MeOH:DCM 1.5:10 v/v) R_F_ = 0.5.

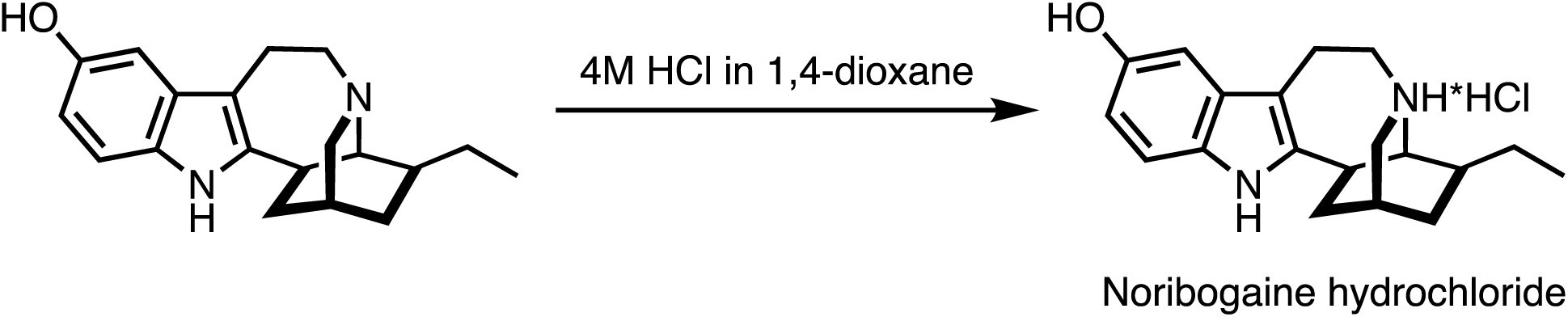

Noribogaine was stirred in 100 mL of 4 M HCl solution in 1,4-dioxane in the dark for 2 hours. The crude was concentrated *in vacuo*, suspended in diethyl ether (100mL), and the precipitate was filtered under reduced pressure. The precipitate was reconstituted in a 30% MeCN in water and lyophilized to yield noribogaine HCl as a beige powder.

**^1^H NMR** (500 MHz, CD_3_OD) δ 7.07 (d, 1H, *J*= 8.5 Hz), 6.79 (d, 1H, *J*= 2.3 Hz), 6.61 (dd, 1H, *J*= 8.5 Hz, 2.3 Hz), 3.55-3.45 (m, 1H), 3.31-3.29 (m, 5H), 3.28-3.14 (m, 5H), 2.21 (t, 1H, *J*= 13.4 Hz), 2.04-1.98 (m, 2H), 1.83 (brS, 1H), 1.71-1.49 (m, 3H), 1.33-1.24 (m, 1H), 0.99 (t, *J*= 7.4 Hz, 3H); **^13^C NMR** (125 MHz, CD_3_OD) δ 149.98, 139.22, 130.10, 128.81, 111.09, 111.05, 105.49, 101.74, 60.15, 56.07, 50.61, 38.84, 34.91, 31.11, 28.71, 25.95, 23.77, 17.97, 10.60; **HRMS** *m/z* (DART): [M+H^+^] calculated for C_19_H_25_ON_2_ 297.4, found 297.2. See Supp. Fig. 6 and Supp. Fig. 7 for ^1^H NMR of noribogaine HCl in CD_3_OD and ^13^C NMR of noribogaine HCl in CD_3_OD.

### Western Blotting

**Table 2.** Western Blotting Protocol Specifics.

| Western | Buffer | Blocking | Primary Antibody |
| --- | --- | --- | --- |
| Fig. 1 Striatum | 20 mM HEPES/1mM EDTA + PI | LI-COR blocking buffer | Goat anti-DAT 1:500, Santa Cruz #sc-1433 (discontinued) |
| Fig. 1 Midbrain | 20 mM HEPES/1mM EDTA + PI | LI-COR blocking buffer | Goat anti-DAT 1:500, Santa Cruz #sc-1433 (discontinued) |
| Fig. 2 | 25mM Tris, 2mM EDTA + PI | 5% BSA in TBS | mouse anti-DAT, 1:250, Thermo Fisher #MA5-24796 |
| Fig. 9 Striata | 30mM phosphate buffer /0.32M sucrose., pH7.9 + PI | 5% BSA in TBS | Rabbit anti-DAT, 1:500, Sigma-Aldrich #D6944 |
| Fig. 9 Midbrains | 30mM phosphate buffer /0.32M sucrose., pH7.9 + PI | 5% BSA in TBS | Rabbit anti-DAT, 1:500, Sigma-Aldrich #D6944 |

BSA, bovine serum albumin; DAT, dopamine transporter; PI, protease inhibitors; TBS, tris-buffered saline

#### Midbrain

For the western blot in Fig. 1, three midbrains per tube were combined and homogenized with glass-teflon homogenizer at setting 7, 12 times in 3 milliliters of 20 mM HEPES/1mM EDTA+ protease inhibitors (PI) leupeptin (Bioshop, LEU001.50), 5 μg/μl pepstatin A (Bioshop, PEP605), 1.5 μg/ml aprotinin (Bioshop, APR600), 0.1 μg/ml benzamidine (Bioshop, BEN601), 100 μm PMSF (Bioshop, PMS123), 2.5 mm Na pyro-phosphate (Bioshop, SPP310), 1 mm β-glycerophosphate (Bioshop, GYP001), 10 mm NaF (Bioshop, SFL001), and 1 mm Na3VO4 (Bioshop SOV664) . The samples were then transferred to a 13 ml tube and homogenized with a polytron for 3×15 seconds at the maximum setting. Samples were then spun at 600g for 10 minutes at 4 degrees. The supernatant was then transferred to a thick walled sorval tube and the pelleted nuclear fractions were discarded. The supernatant was spun at 40,000g for 15 minutes at 4 degrees. The supernatant was then discarded and the pellet was resuspended in 100ul of HEPES/EDTA+PI buffer. Protein concentration was determined using a BCA protein assay kit (Thermo Fisher Scientific, Pierce BCA Protein Assay Kit, #23227). Membrane protein extracts were then separated by 10% SDS-PAGE and transferred to a PVDF membrane. Approximately even loading and transfer were confirmed using Ponceau S stain before proceeding with immunodetection. Blot was blocked for 1 hour at room temperature with LI-COR (Lincoln, NE) blocking buffer and stained overnight at 4 degrees with goat anti-DAT (1:500, Santa Cruz #sc-1433 [discontinued]). The blot was then incubated with a donkey anti-goat secondary for 1 hour at room temperature (1:10000). Protein bands were visualized using the Odyssey Imaging System (LI-COR) and Image Studio software.

For the midbrain western blots in Fig. 12, individual midbrains were homogenized in 2mL of 30mM phosphate buffer (pH 7.9) containing 0.32M sucrose + PIs. Homogenates were centrifuged at 1000 rpm at 4 degrees for 10 minutes. The supernatant was transferred to a new tube and centrifuged twice at 18000 rpm for 20 minutes at 4 degrees Celsius. The supernatant was discarded and the resulting pellet was resuspended in 100ul of buffer. Protein concentration was determined using a BCA protein assay kit (Thermo Fisher Scientific, Pierce BCA Protein Assay Kit, #23227). Membrane protein extracts were then separated by 10% SDS-PAGE and transferred to a PVDF membrane. Approximately even loading and transfer were confirmed using LI-COR Total Protein stain (LI-COR, LIC-926-11010) before proceeding with immunodetection. Blots were blocked for 1 hour at room temperature with LI-COR (Lincoln, NE) blocking buffer and stained overnight at 4 degrees with a rabbit anti-DAT primary antibody (1:500, Sigma-Aldrich, #D6944). The blots were then incubated with an anti-rabbit secondary for 1 hour at room temperature (1:7500). Protein bands were visualized using the Odyssey Imaging System (LI-COR) and Emperia Studio software. Levels of DAT were calculated as follows: ((DAT signal/Total Protein Signal)/(Vehicle Average (DAT signal/Total Protein Signal)) * 100.

#### Striatum

Striata in Fig. 1 were homogenized with glass-teflon homogenizer at setting 7, 12 times in 3ml of 20 mM HEPES/1mM EDTA or 25 mm Tris-2 mM EDTA, and 10 μg/ml of PIs leupeptin (Bioshop, LEU001.50), 5 μg/μl pepstatin A (Bioshop, PEP605), 1.5 μg/ml aprotinin (Bioshop, APR600), 0.1 μg/ml benzamidine (Bioshop, BEN601), 100 μm PMSF (Bioshop, PMS123), 2.5 mm Na pyro-phosphate (Bioshop, SPP310), 1 mm β-glycerophosphate (Bioshop, GYP001), 10 mm NaF (Bioshop, SFL001), and 1 mm Na3VO4 (Bioshop SOV664). The samples were then transferred to a 13 ml tube and homogenized with a polytron for 3×15 seconds at the maximum setting. Samples were then spun at 600g for 10 minutes at 4 degrees. The supernatant was then transferred to a thick walled sorval tube and the pelleted nuclear fractions were discarded. The supernatant was spun at 40,000g for 15 minutes at 4 degrees. The supernatant was then discarded and the pellet was resuspended in 100ul of HEPES/EDTA+PI buffer. Protein concentration was determined using a BCA protein assay kit (Thermo Fischer Scientific, Pierce BCA Protein Assay Kit, #23227). Membrane protein extracts were then separated by 10% SDS-PAGE and transferred to a PVDF membrane. Approximately even loading and transfer were confirmed using Ponceau S stain before proceeding with immunodetection. Blot was blocked for 1 hour at room temperature with LI-COR (Lincoln, NE) blocking buffer and stained overnight at 4 degrees with goat anti-DAT (1:500, Santa Cruz #sc-1433 [discontinued]). The blot was then incubated with a donkey anti-goat secondary for 1 hour at room temperature (1:10000) overnight at 4 degrees Celsius. Protein bands were visualized using the Odyssey Imaging System (LI-COR) and Image Studio software or Emperia Studio software. . Levels of DAT were calculated as follows: ((DAT signal/Total Protein Signal)/(Vehicle Average (DAT signal/Total Protein Signal)) * 100.

Striata in Fig. 12 were homogenized in 2mL of 30mM phosphate buffer (pH 7.9) containing 0.32M sucrose + PIs. Homogenates were centrifuged at 1000 rpm at 4 degrees for 10 minutes. The supernatant was transferred to a new tube and centrifuged twice at 18000 rpm for 20 minutes at 4 degrees Celsius. The supernatant was discarded and the resulting pellet was resuspended in 100ul of buffer. Protein concentration was determined using a BCA protein assay kit (Thermo Fisher Scientific, Pierce BCA Protein Assay Kit, #23227). Membrane protein extracts were then separated by 10% SDS-PAGE and transferred to a PVDF membrane. Approximately even loading and transfer were confirmed using LI-COR Total Protein stain (LI-COR, LIC-926-11010) before proceeding with immunodetection. Blot was blocked for 1 hour at room temperature with 5% BSA in TBS and stained overnight at 4 degrees with rabbit anti-DAT primary antibody (1:500, Sigma-Aldrich, #D6944). The blot was then incubated with an anti-rabbit secondary for 1 hour at room temperature (1:7500). Protein bands were visualized using the Odyssey Imaging System (LI-COR) and Emperia Studio software. Levels of DAT were calculated as follows: ((DAT signal/Total Protein Signal)/(Vehicle Average (DAT signal/Total Protein Signal)) * 100.

#### EndoH/PNGase Experiment

The midbrain (3 samples/tube) was dissected and homogenized in a buffer containing 25mM Tris, 2mM EDTA, 10ug/ml leupeptin, 5ug/ml pepstatin A, 1.5ug/ml aprotinin, 0.1ug/ml benzamidine, 100uMPMSF, 2.5mMNaPyro, 1mM beta-glycerophosphate, 10mMNaF, and 1mMNaVO4. The samples were incubated on ice for 10 minutes and spun at 13,000 rpm for 10 minutes at 4 degrees. Protein concentrations were determined using a BCA protein assay kit (Thermo Fisher Scientific, Pierce BCA Protein Assay Kit, #23227). Samples were digested with the glycosidases peptide N-glycosidase F (PNGase F) and EndoH (New England Biolabs, catalog #P0704S and #P0702S, respectively). Membrane protein extracts were then separated by 10% SDS-PAGE and transferred to PVDF membranes. Approximately even loading and transfer were confirmed using LI-COR Total Protein stain (LI-COR, LIC-926-11010) before proceeding with immunodetection. Blots were blocked with 5% BSA in TBS for 1 hour at room temperature and stained for 1 hour at room temperature and then overnight at 4 degrees Celsius with mouse anti-DAT (1:250, Thermo Fisher Scientific, MA, USA, #MA5-24796). The blots were then incubated with a secondary anti-mouse (1:7500) antibody. Protein bands were visualized using the Odyssey Imaging System (LI-COR) and Image Studio software.

#### Tissue Content of Dopamine and Metabolites

Dopamine levels were determined using high-performance liquid chromatography with electrochemical detection (HPLC-EC; ESA/Thermo Scientific, Chelmsford, MA; 37). Separation of neurotransmitters and their metabolites was achieved using a reverse phase column (Luna 100 × 3.0 mm C18 3 μm, Phenomenex, Torrance, CA) with a mobile phase made up of 75 mM NaH2PO4, 1.7 mM 1-octanesulfonic acid sodium salt, 100 µL/L triethylamine, 25 µM EDTA, 10% acetonitrile v/v; pH 3.0. Analyte detection was carried out using a high-sensitivity analytical cell 5011 A (Thermo Fisher Scientific, Sunnyvale, CA) at +220 mV on a Coulochem III Electrochemical Detector (ESA/Thermo Scientific, Chelmsford, MA). Chromeleon software (ESA/Thermo Scientific, Chelmsford, MA) was used to quantify analytes using an external calibration curve of standards of known dopamine and metabolite concentrations.

#### Fast-Scan Cyclic Voltammetry (FSCV)

Mice were anesthetized with 5% isoflurane and decapitated. Brains were removed and placed into oxygenated (95% O2 / 5% CO2) artificial cerebrospinal fluid composed of (in mM): 126 mM NaCl, 2.5 mM KCl, 1.2 mM NaH2PO4, 2.4 mM CaCl2, 1.2 mM MgCl2, 25 mM NaHCO3, 11mM D-glucose, and 0.4 mM L-ascorbic acid, with pH adjusted to 7.4. Brains were sliced into 300 µm coronal sections between +1.7 to +0.74 mm from bregma with a vibrating tissue slicer. Slices were allowed to equilibrate for one hour in oxygenated aCSF flowing at 1 mL/min. A glass capillary-pulled carbon fiber recording microelectrode was placed 75 µm into the slice next to a bipolar stimulating electrode. Dopamine release was evoked by single monophasic electrical pulses (750 µA, 4 ms) occurring every 300 seconds until signals were stable (<10% change in peak height) for at least three measurements. Recordings were taken for 15 seconds (wild-type) or 30 seconds (A313V) by the recording electrode that scanned a triangular waveform between -0.4 V and 1.2 V against Ag/AgCl reference at a rate of 400 V/s every 100 ms. Dopamine oxidation current (nA) was converted to concentration (µM) with calibration factors based on the size of the electrode background current, which reflects active carbon surface area and thereby sensitivity to dopamine, to predict the magnitude of signal in response to a known (3 µM) concentration of dopamine, based on 100+ typical electrode responses. Dopamine recordings were taken in the dorsolateral striatum. Recording and analysis were conducted with Demon Voltammetry and Analysis Software (Yorgason et al., 2011). Stable baseline collections were first analyzed using a least squares fitting equation to assess peak height and decay (tau). Subsequently, baseline recordings and drug collections were analyzed via Michaelis-Menten kinetic modeling to determine dopamine release (µM) and maximal rate of dopamine uptake (Vmax; µM/s). Representative traces were generated by taking the raw current versus time plots for one stable baseline recording per slice, then dividing all time points in that recording by the electrode calibration factor for that experiment.

### Behavioral assessments

#### Dorsal Immobility Response

Animals were clasped by the nape of the neck and gently elevated into the air to elicit the dorsal immobility response. Latency to begin moving was recorded for each animal.

#### Open Field Test

Locomotor activity and stereotypy were measured using digital activity monitors (Omnitech Electronics, Columbus, OH, USA). Animals were placed in Plexiglas chambers (20 × 20 × 45 cm3) and their locomotor activity and stereotypic behavior were recorded. After 60 minutes, mice were given i.p. injections of vehicle or drug. Infrared light beam sensors were used to track the animal’s movement. Total distance traveled in centimeters, horizontal activity, vertical activity, stereotypy number and stereotypy time were collected in 5-min bins. Mice were naïve to the task and had not been exposed to the arena prior to testing. Data was discarded for mice who escaped from the arena during testing. Data were combined for animals that received 0.9% saline as a drug vehicle and whose activity was monitored using the VersaMax Legacy Open Field equipment (Omnitech Electronics, Inc. Columbus, Ohio, USA) (baseline, 30 mg/kg MPH, 40 mg/kg MPH, 250 mg/kg αMPT, 125 mg/kg αMPT. and 62.5 m/kg αMPT. Open field experiments performed using the VersaMax SuperFlex Open Field System and Fusion Software (Omnitech Electronics, Inc. Columbus, Ohio, USA) had their own dedicated vehicle groups and were not combined with others (31 mg/kg αMPT, noribogaine).

#### Habituation Index

The habituation index, representing the time in minutes to reach half-maximal activity in the open field test, was calculated following Mielnik et al., 2021. To determine this index, the Y-intercept from the activity data between the first time point (minute 5) and the last time point (minute 150) was divided by 2, and this result was then divided by the slope of the data from minute 5 to minute 150. The absolute value of this calculation was used for statistical testing for each animal.

#### Rotarod

A computer controlled rotarod apparatus (46750 Rota-Rod NG, Ugo Basile, Gemonio, VA, Italy) with a rod of 3 cm diameter was set to accelerate from 4 to 40 revolutions per minute over 300 seconds, and latency to fall was recorded. Motor learning was assessed using a method adapted from Beeler et al., 2010, wherein mice received 5 consecutive trials per session, 1 session per day. Rest between trials was approximately 30 seconds. Mice were trained for 5 days, then received a 3 day break, and were tested again on the 9th day.

#### Gait Analysis

Gait analysis was performed using the DigiGait treadmill by Mouse Specifics, Inc. The equipment consists of a walking compartment of 25cm with a clear polymer bonded belt. A high-speed digital video camera images the underside of the walking animals. Gait was recorded from each animal for a minimum of 3 seconds at a walking speed of 20 centimeters per second. The DigiGait software generated digital paw prints and dynamic gait signals that were analyzed automatically using MatLab. The positions of 14 body parts were labeled in each video frame (nose, tail base and tip, torso center, left and right torso flanks, base of palm and 1st joint of middle digit on each paw) using custom neural network models in the Python 3.11-based DeepLabCut v2.3.7 software toolbox (see deeplabcut.github.io/DeepLabCut/; Nath et al., 2019). Two Resnet-50-based network models were trained to estimate the position of an animal’s 4 paws either A) throughout all video frames, or B) exclusively when a given paw was planted on the treadmill surface. The latter model (accessible at osf.io/d3wc6) was used to extract the stride length dataset reported here. Asymptotic model performance (2.24 px train error and 2.55 px test error with a 0.6 probability estimate cutoff) was reached following 150000 training iterations, including manual labeling a total of 340 video frames from 16 training set video segments. Tail tip was inconsistently visible throughout each video and did not contribute to subsequent analyses. An experimenter blinded to experimental design/group manually corrected any residual tracking errors while viewing DLC-labeled body parts overlaid on each video snippet (see tracking example video at https://osf.io/gm6hs/).

In total, 5 complete stride cycles were analyzed for each experimental subject (5 left and 5 right strides), curated based on the visibility of all body parts during continuous forward walking motion (excluding intervals in which the mouse displayed prominent lateral locomotion, grooming, rearing, ground/wall whisking, halted momentum). These exclusion criteria were implemented to isolate differences in fine motor coordination during forward locomotion independently from gross differences in behavioral activity (i.e. exploration, arousal/excitability, compulsive grooming, etc.), which are better captured in the OFT and other behavioral assays. Stride length was calculated as the distance between the forward- and backward-most extension, respectively, of the forepaw and hind paw on each side of the body. As these positions typically occurred asynchronously during a stride, nose position was used to correct for overall changes in body position across frames (thus isolating stride length relative to the mouse, despite ongoing acceleration/deceleration on the treadmill). Animal length (mean difference in Cartesian position of nose and tail base in each frame that met inclusion criteria) was used to scale the stride length as a proportion of body length. The mean body lengths of WT (398.89 px) and Het (399.34 px) groups did not differ statistically (t42 = 0.0898, p = 0.929). Thus, the mean stride length for each animal was calculated as the average of 10 strides, as a percentage of their body length. Microsoft Excel was used to compute all relevant position metrics from raw DLC pose estimates.

### Noribogaine experiments

Noribogaine was suspended in 5% ethanol, 5% glucose at a concentration of 10, 25, or 50 mg/mL for doses of 100 mg/kg, 250 mg/kg, or 500 mg/kg, respectively. Noribogaine was administered via oral gavage. Blood was collected as indicated from the saphenous vein using lithium heparin tubes (Sarstedt #16.443.100). Blood samples were spun at 5,000*g* for 10 minutes, and the supernatant (plasma) was collected for further analysis. Animals were killed via rapid conscious cervical dislocation and brains were flash frozen in isopentane. Midbrains and striata were dissected for analysis of DAT levels via western blotting, and cerebellums were dissected for noribogaine concentration analysis via LCMS.

### LC-MS/MSNoribogaine Plasma and Brain Concentration Analysis

Concentrations of noribogaine in plasma and brain were measured using LC-MS/MS with clonazepam-d_4_ (CLZ-d_4_) as internal standard. The analysis was done on an Agilent 1260 LC system (Agilent 1260 Quaternary pump, Agilent 1260 Infinity Standard Autosampler and temperature-controlled column compartment) using a C18 column (Zorbax SB-C18, 5µm 2.1 x 150 mm, 5µm; Agilent) at ambient temperature. Noribogaine and internal standard were resolved with a gradient elution where the mobile phase A was Milli-Q water + 0.1% formic acid, and B was HPLC-grade acetonitrile + 0.1% formic acid. Gradient elution was used as the following:

**Table 3.** LCMS gradient elution.

| Time (min) | Flow (ml/min) | (A) | (B) |
| --- | --- | --- | --- |
| 0 | 0.40 | 90 | 10 |
| 1.0 | 0.40 | 90 | 10 |
| 10.0 | 0.40 | 40 | 60 |
| 11.0 | 0.40 | 90 | 10 |
| 15 | 0.40 | 90 | 10 |

The LC system was coupled to an Agilent 6430 triple quadropole mass spectrometer (Agilent,Santa Clara, CA) operated in positive Electrospray Ionization (ESI) mode with source temperature maintained at 350°C. The mass transitions monitored were m/z 297.2 to 122.1 (used for quantification) and 297.2 to160 for noribogaine and 320.1 to 274 for CLZ-d_4._

For sample preparation, a stock mixture of the internal standard solution of CLZ-d4 was dissolved in methanol with 1% formic acid to a concentration of 50 ng/mL. Calibration curves were prepared in plasma for analysis of plasma samples (1, 10, 100, 500, 1000 and 5000 ng/mL) and in brain homogenate where 1 part tissue was homogenized in 3 volumes of 0.01 HCl (1, 10, 100, 500, 1000 and 5000 ng/mL). For analysis of the plasma samples, 40 µL of 50 ng/mL internal standard solution was added to 20µL of the plasma sample or calibrator for a final concentration of 100 ng/mL in plasma. Samples and calibrators were vortexed vigorously for mixing, and centrifuged at approximately 15,000 g (12,000 rpm on small tabletop centrifuge) for 15 minutes,30 µL of the supernatant was transferred to LCMS vial inserts and 30 µL of water was added to reduce methanol content. The injection volume used was 20 µL. For the brain samples, 100 µL of 50 ng/mL internal standard was added to 50 µL of the brain homogenate (1:3 in 0.01 M HCl) sample or calibrator for a final concentration of 100 ng/mL in brain homogenate. Samples and calibrators were vortexed vigorously for mixing and centrifuged at approximately ∼ 15,000 g (12,000 rpm on small tabletop centrifuge) for 15 minutes,50 µL of the supernatant was transferred to LCMS vial inserts and 50 µL of water was added to reduce methanol content. The injection volume was 20 µL.

Recovery and matrix effects were determined for two concentrations (200 and 2,000 ng/mL) for plasma and brain homogenate to range between 71.1-86.8% for noribogaine in brain homogenate and 37.1-49.4% in plasma, respectively. The internal standard peak area response ratio was tested for linearity between the concentration 10 to 10,000 ng/ml). Six calibration samples are enough to describe the relationship (10, 100, 500, 1,000, 5,000 and 10,000 ng/ml) with weighing (1/x2). The limit of quantification (LOQ) was determined to 10 ng/mL for noribogaine. Intraday accuracy and precision were determined using replicates of 3 QC samples (10, 200 and 2000 ng/ml) in plasma and brain homogenate matrixes. . Inter-day accuracy was 85.4 - 113.4% for plasma and 90.9 - 112.3% for brain homogenate, and precision (%CV) was 8.8-10.2% for plasma and 5.1 - 11.2% for brain homogenate.

### Statistical Analyses

Outliers within each group were identified and removed using the interquartile range method. For each variable, the first quartile (Q1) and third quartile (Q3) were calculated. The interquartile range (IQR) was defined as the difference between Q3 and Q1 (IQR=Q3-Q1). Data points were considered outliers if they fell below the lower bound (Q1-1.5 x IQR) or above the upper bound (Q3 + 1.5 × IQR). Statistics were performed after outliers were removed. All statistical analyses were performed for each group by a Student’s t-test , two-way ANOVA, or mixed model, indicated in each figure legend where appropriate. Tukey’s or Šidák’s multiple comparisons were performed after the ANOVA or mixed model analysis if the interaction between variables was significant. Significance is reported where p≤0.05. All statistical testing was carried out using GraphPad Prism version 9 or 10.

### Study Approval

Animal housing and experiments were conducted in accordance with the Canadian Council on Animal Care and the University of Toronto Faculty of Medicine and Pharmacy Animal Care Committee.

## Acknowledgements

We would like to acknowledge Dr. Iain Greig, Wendy Horsfall, Lola Zovko, Dr. Yuanye (Phoebe) Yan, Megan Sullivan, Sylvia Okafor, Lyra Vania, Dr. Ilias Marmouzi and all members of Ramsey and Salahpour labs for helpful discussions.

## Author Contributions

E.E.R., A.R., C.W., P.B., T.V.L., P.S.B.F., L.J.E., R.F.T., D.W.C., A.J.R., S.R.J., and A.S. designed research; E.E.R., A.R., C.W., E.Q.W., P.B., M.M., M.N., A.B., T.V.L., J.L., R.C., P.S.B.F., and D.W.C performed research; E.E.R., A.R., C.W., P.B., M.M., M.N., A.B., T.V.L., J.L., R.C., P.S.B.F., R.F.T., D.W.C., A.J.R., S.R.J., and A.S. analyzed data; E.E.R., and A.S. wrote the paper; all authors were involved in editing the manuscript.

## References

1. Ciliax, B. J et al., (1995). The dopamine transporter: Immunochemical characterization and localization in brain. J Neurosci, 15(3 Pt 1), 1714–1723.

2. Ciliax, B. J et al., (1999). Immunocytochemical localization of the dopamine transporter in human brain. J Comp Neurol, 409(1), 38–56. 10.1002/(sici)1096-9861(19990621)409:1<38::aid-cne4>3.0.co;2-1

3. Chen, N., & Reith, M. E. (2000). Structure and function of the dopamine transporter. Eur J Pharmacol, 405(1–3), 329–339. 10.1016/s0014-2999(00)00563-x

4. Bu, M., Farrer, M. J., & Khoshbouei, H. (2021). Dynamic control of the dopamine transporter in neurotransmission and homeostasis. NPJ Parkinson’s Dis, 7(1), 1–11. 10.1038/s41531-021-00161-2

5. Giros, B. et al., (1996). Hyperlocomotion and indifference to cocaine and amphetamine in mice lacking the dopamine transporter. Nature, 379(6566), 606–612. 10.1038/379606a0

6. Salahpour, A. et al., (2008). Increased amphetamine-induced hyperactivity and reward in mice overexpressing the dopamine transporter. Proc Natl Acad Sci U S A, 105(11), 4405– 4410. 10.1073/pnas.0707646105

7. Vaughan, R. A., & Foster, J. D. (2013). Mechanisms of dopamine transporter regulation in normal and disease states. Trends Pharmacol Sci, 34(9), 489–496. 10.1016/j.tips.2013.07.005

8. Kanner, B. I., & Zomot, E. (2008). Sodium-coupled neurotransmitter transporters. Chem Rev, 108(5), 1654–1668. 10.1021/cr078246a

9. Vaughan, R. A., & Foster, J. D. (2013). Mechanisms of dopamine transporter regulation in normal and disease states. Trends Pharmacol Sci, 34(9), 489–496. 10.1016/j.tips.2013.07.005

10. DiCarlo, G. E. et al., (2019). Autism-linked dopamine transporter mutation alters striatal dopamine neurotransmission and dopamine-dependent behaviors. Journal Clin Invest, 129(8), 3407–3419. 10.1172/JCI127411

11. Hamilton, P. J. et al., (2013). De novo mutation in the dopamine transporter gene associates dopamine dysfunction with autism spectrum disorder. Mol Psychiatry, 18(12), 1315–1323. 10.1038/mp.2013.102

12. Mazei-Robison, M. S. et al., (2005). Sequence variation in the human dopamine transporter gene in children with attention deficit hyperactivity disorder. Neuropharmacology, 49(6), 724–736. 10.1016/j.neuropharm.2005.08.003

13. Pinsonneault, J. K., et al., (2011). Dopamine Transporter Gene Variant Affecting Expression in Human Brain is Associated with Bipolar Disorder. Neuropsychopharmacology, 36(8), 1644–1655. 10.1038/npp.2011.45

14. Assmann, B. et al., (2004). Infantile parkinsonism-dystonia and elevated dopamine metabolites in CSF. Neurology, 62(10), 1872–1874. 10.1212/01.WNL.0000126440.16612.51

15. Kurian, M. A. et al., (2009). Homozygous loss-of-function mutations in the gene encoding the dopamine transporter are associated with infantile parkinsonism-dystonia. J Clinl Invest, 119(6), 1595–1603. 10.1172/JCI39060

16. Kurian, M. A. et al., (2011). Clinical and molecular characterisation of hereditary dopamine transporter deficiency syndrome: An observational cohort and experimental study. Lancet Neurol, 10(1), 54–62. 10.1016/S1474-4422(10)70269-6

17. Ng, J. et al., (2014). Dopamine transporter deficiency syndrome: Phenotypic spectrum from infancy to adulthood. Brain, 137(4), 1107–1119. 10.1093/brain/awu022

18. Yildiz, Y. et al., (2016). Hereditary Dopamine Transporter Deficiency Syndrome: Challenges in Diagnosis and Treatment. Neuropediatrics, 48, 49–52. 10.1055/s-0036-1593372

19. Galiart, E. et al., (2017). Infantile Dystonia Parkinsonism Caused by a Mutation in SLC6A3: Case Report of Three Siblings. Neuropediatrics, 48, VS06. 10.1055/s-0037-1603021

20. Kuster, A. et al., (2018). Diagnostic approach to neurotransmitter monoamine disorders: Experience from clinical, biochemical, and genetic profiles. J Inherit Metab Dis, 41(1), 129–139. 10.1007/s10545-017-0079-6

21. Heidari, E. et al., (2020). Homozygous in-frame variant of SCL6A3 causes dopamine transporter deficiency syndrome in a consanguineous family. Ann of Hum Genet, 84(4), 315–323. 10.1111/ahg.12378

22. Nasehi, M. M. et al., (2020). Dopamine Transporter Deficiency Syndrome: A Case with Hyper- and Hypokinetic Extremes. Mov Disord Clin Pract, 7(Suppl 3), S57–S60. 10.1002/mdc3.13064

23. Tehreem, B., & Kornitzer, J. (2020). Expanding the phenotypic spectrum of dopamine transporter deficiency syndrome with a novel mutation (1884). Neurology, 94(15_supplement), 1884. 10.1212/WNL.94.15_supplement.1884

24. Baga, M., Spagnoli, C., Soliani, L., Salerno, G. G., Rizzi, S., Frattini, D., Pisani, F., & Fusco, C. (2021). Early-onset Dopamine Transporter Deficiency Syndrome: Long-term Follow-up. Can J Neurol Sci, 48(2), 285–286. 10.1017/cjn.2020.144

25. Mir, A. et al., (2022). SLC gene mutations and pediatric neurological disorders: diverse clinical phenotypes in a Saudi Arabian population. Hum Genet 141, 81–99 (2022).

26. Silva, A. P. R. et al., (2023). Case report of two brothers with infantile Parkinsonism-dystonia (OMIM #613135). Arq Neuropsiquiatr, 81, A226. 10.1055/s-0043-1774645

27. Spaull, R. V., & Kurian, M. A. (1993). SLC6A3-Related Dopamine Transporter Deficiency Syndrome. In M. P. Adam, J. Feldman, G. M. Mirzaa, R. A. Pagon, S. E. Wallace, & A. Amemiya (Eds.), GeneReviews®. University of Washington, Seattle. http://www.ncbi.nlm.nih.gov/books/NBK442323/

28. Sanders, C. R., & Myers, J. K. (2004). Disease-related misassembly of membrane proteins. Annu Rev Biophys Biomol Struct, 33, 25–51. 10.1146/annurev.biophys.33.110502.140348

29. Morello, J. P. et al., (2000). Pharmacological chaperones: A new twist on receptor folding. Trends Pharmacol Sci, 21(12), 466–469. 10.1016/s0165-6147(00)01575-3

30. Beerepoot, P. et al., (2016). Pharmacological Chaperones of the Dopamine Transporter Rescue Dopamine Transporter Deficiency Syndrome Mutations in Heterologous Cells. Biol Chem, 291(42), 22053–22062. 10.1074/jbc.M116.749119

31. Asjad, H. M. M. et al., (2017). Pharmacochaperoning in a Drosophila model system rescues human dopamine transporter variants associated with infantile/juvenile parkinsonism. Biol Chem, 292(47), 19250–19265. 10.1074/jbc.M117.797092

32. Leidenheimer, N. J., & Ryder, K. G. (2014). Pharmacological Chaperoning: A Primer on Mechanism and Pharmacology. Pharmacol Res, 83, 10–19. 10.1016/j.phrs.2014.01.005

33. Bhat, S. et al., (2021). Tropane-Based Ibogaine Analog Rescues Folding-Deficient Serotonin and Dopamine Transporters. ACS Pharmacol Transl Sci, 4(2), 503–516. 10.1021/acsptsci.0c00102

34. Morello, J. P. et al., (2000). Pharmacological chaperones: A new twist on receptor folding. Trends Pharmacol Sci 21(12), 466–469. 10.1016/s0165-6147(00)01575-3

35. Germain, D. P., et al., (2019). Efficacy of the pharmacologic chaperone migalastat in a subset of male patients with the classic phenotype of Fabry disease and migalastat-amenable variants: Data from the phase 3 randomized, multicenter, double-blind clinical trial and extension study. Genet Med, 21(9), 1987–1997. 10.1038/s41436-019-0451-z

36. Baatallah, N. et al., (2021). Pharmacological chaperones improve intra-domain stability and inter-domain assembly via distinct binding sites to rescue misfolded CFTR. Cell Mol Life Sci, 78(23), 7813–7829. 10.1007/s00018-021-03994-5

37. Janovick, J. A. et al., (2013). Restoration of testis function in hypogonadotropic hypogonadal mice harboring a misfolded GnRHR mutant by pharmacoperone drug therapy. Proc Natl Acad Sci U S A, 110(52), 21030–21035. 10.1073/pnas.1315194110

38. Freissmuth, M. et al., (2018). SLC6 Transporter Folding Diseases and Pharmacochaperoning. Handb Exp Pharmacol, 245, 249–270. 10.1007/164_2017_71

39. El-Kasaby, A. et al., (2024). Allosteric Inhibition and Pharmacochaperoning of the Serotonin Transporter by the Antidepressant Drugs Trazodone and Nefazodone. Mol Pharmacol, 106(1), 56–70. 10.1124/molpharm.124.000881

40. El-Kasaby, A. et al., (2010). Mutations in the Carboxyl-terminal SEC24 Binding Motif of the Serotonin Transporter Impair Folding of the Transporter*. Biol Chem, 285(50), 39201–39210. 10.1074/jbc.M110.118000

41. Kasture, A. et al., (2016). Functional Rescue of a Misfolded Drosophila melanogaster Dopamine Transporter Mutant Associated with a Sleepless Phenotype by Pharmacological Chaperones *♦. Biol Chem, 291(40), 20876–20890. 10.1074/jbc.M116.737551

42. Litjens, R. P. W., & Brunt, T. M. (2016). How toxic is ibogaine? Clin Toxicol, 54(4), 297–302. 10.3109/15563650.2016.1138226

43. Sutton, C. et al., (2022). Structure-Activity Relationships of Dopamine Transporter Pharmacological Chaperones. Front Cell Neurosci, 16. 10.3389/fncel.2022.832536

44. Jones, S. R. et al., (1998). Profound neuronal plasticity in response to inactivation of the dopamine transporter. Proc Natl Acad Sci U S A, 95(7), 4029–4034. 10.1073/pnas.95.7.4029

45. Sotnikova, T. D. et al., (2005). Dopamine-Independent Locomotor Actions of Amphetamines in a Novel Acute Mouse Model of Parkinson Disease. PLoS Biol, 3(8), e271. 10.1371/journal.pbio.0030271

46. Bossé, R. et al., (1997). Anterior pituitary hypoplasia and dwarfism in mice lacking the dopamine transporter. Neuron, 19(1), 127–138. 10.1016/s0896-6273(00)80353-0

47. Lalonde, R., & Strazielle, C. (2022). Neurochemical anatomy of dorsal and tonic immobility responses. Pharmacol Biochem and Behav, 213, 173334. 10.1016/j.pbb.2022.173334

48. Mielnik, C. A. et al.,(2021). Consequences of NMDA receptor deficiency can be rescued in the adult brain. Mol Psychiatry, 26(7), 2929–2942. 10.1038/s41380-020-00859-4

49. Cyr, M. et al., (2003). Sustained elevation of extracellular dopamine causes motor dysfunction and selective degeneration of striatal GABAergic neurons. Proc Natl Acad Sci U S A, 100(19), 11035–11040. 10.1073/pnas.1831768100

50. Ogura, T. et al., (2005). Impaired acquisition of skilled behavior in rotarod task by moderate depletion of striatal dopamine in a pre-symptomatic stage model of Parkinson’s disease. Neurosci Res, 51(3), 299–308. 10.1016/j.neures.2004.12.006

51. Beeler, J. A. et al., (2010). Dopamine-dependent motor learning: Insight into levodopa’s long-duration response. Ann Neurol, 67(5), 639–647. 10.1002/ana.21947

52. Aguilar, J. I. et al., (2021). Psychomotor impairments and therapeutic implications revealed by a mutation associated with infantile Parkinsonism-Dystonia. eLife, 10, e68039. 10.7554/eLife.68039

53. Deacon, R. M. J. (2013). Measuring Motor Coordination in Mice. Journal Vis Exp, 75, e2609. 10.3791/2609

54. Gainetdinov, R. R. et al., (1999). Role of serotonin in the paradoxical calming effect of psychostimulants on hyperactivity. Sci, 283(5400), 397–401. 10.1126/science.283.5400.397

55. Sotnikova, T. D. et al., (2005). Dopamine-Independent Locomotor Actions of Amphetamines in a Novel Acute Mouse Model of Parkinson Disease. PLoS Biol, 3(8), e271. 10.1371/journal.pbio.0030271

56. Mash, D. C. et al., (2016). Oral noribogaine shows high brain uptake and anti-withdrawal effects not associated with place preference in rodents. J Psychopharmacol, 30(7), 688– 697. 10.1177/0269881116641331

57. Schubert, U. et al.,(2000). Rapid degradation of a large fraction of newly synthesized proteins by proteasomes. Nature, 404(6779), 770–774. 10.1038/35008096

58. Guerois, R. et al., (2002). Predicting Changes in the Stability of Proteins and Protein Complexes: A Study of More Than 1000 Mutations. J Mol Biol, 320(2), 369–387. 10.1016/S0022-2836(02)00442-4

59. Vecchio, L. M. et al., (2014). N-terminal tagging of the dopamine transporter impairs protein expression and trafficking in vivo. Mol Cell Neurosci, 61, 123–132. 10.1016/j.mcn.2014.05.007

60. Nepal, B. et al., (2023). Overview of the structure and function of the dopamine transporter and its protein interactions. Front Physiol, 14. 10.3389/fphys.2023.1150355

61. Eisenhofer, G. et al., (2004). Catecholamine metabolism: A contemporary view with implications for physiology and medicine. Pharmacol Rev, 56(3), 331–349. 10.1124/pr.56.3.1

62. Meiser, J. et al., (2013). Complexity of dopamine metabolism. Cell Commun Signal, 11(1), 34. 10.1186/1478-811X-11-34

63. Kaufman, D. M. Neurotransmitters and Drug Abuse. In: Kaufman, D. M., ed. Clinical Neurology for Psychiatrists (Sixth Edition). W.B. Saunders; 2007:11–536. 10.1016/B978-1-4160-3074-4.10021-9

64. Myöhänen, T. T. et al., (2010). Distribution of catechol-O-methyltransferase (COMT) proteins and enzymatic activities in wild-type and soluble COMT deficient mice. J Neurochem, 113(6), 1632–1643. 10.1111/j.1471-4159.2010.06723.x

65. Holleran, K. M. et al., (2020). Organic cation transporter 3 and the dopamine transporter differentially regulate catecholamine uptake in the basolateral amygdala and nucleus accumbens. Eur J Neurosci, 10.1111/ejn.14927.

66. Martin, D., & Le, J. K. (2025). Amphetamine. In StatPearls. StatPearls Publishing. http://www.ncbi.nlm.nih.gov/books/NBK556103/

67. Brogden, R.N., et al., (1981). α-Methyl-*p*-Tyrosine: A Review of its Pharmacology and Clinical Use. Drugs 21, 81–89. https://doiorg.myaccess.library.utoronto.ca/10.2165/00003495-198121020-00001

68. Hattori, A. et al., (1998). Intrastriatal injection of dopamine results in DNA damage and apoptosis in rats. NeuroReport, 9(11), 2569.

69. Liu, Z. et al., (2001). A novel mechanism of dopamine neurotoxicity involving the peripheral extracellular and the plasma membrane dopamine transporter. NeuroReport, 12(15), 3293.

70. Chen, L. et al., (2008). Unregulated Cytosolic Dopamine Causes Neurodegeneration Associated with Oxidative Stress in Mice. J Neurosci, 28(2), 425–433. 10.1523/JNEUROSCI.3602-07.2008

71. Jiang, Y. et al., (2008). Extracellular dopamine induces the oxidative toxicity of SH-SY5Y cells. Synapse, 62(11), 797–803. 10.1002/syn.20554

72. Mosharov, E. V. et al., (2009). Interplay between cytosolic dopamine, calcium, and alpha-synuclein causes selective death of substantia nigra neurons. Neuron, 62(2), 218–229. 10.1016/j.neuron.2009.01.033

73. Benedetto, A. et al., (2010). Extracellular Dopamine Potentiates Mn-Induced Oxidative Stress, Lifespan Reduction, and Dopaminergic Neurodegeneration in a BLI-3–Dependent Manner in Caenorhabditis elegans. PLOS Genet, 6(8), e1001084. 10.1371/journal.pgen.1001084

74. Young, William F. Endocrine Hypertension. In: S. Melmed, K. S. Polonsky, P. R. Larsen, & H. M. Kronenberg, eds. Williams Textbook of Endocrinology (Thirteenth Edition). Elsevier; 2016:556–588. 10.1016/B978-0-323-29738-7.00016-2

75. Golden, J. P. et al., (2013). Dopamine-Dependent Compensation Maintains Motor Behavior in Mice with Developmental Ablation of Dopaminergic Neurons. J Neurosci, 33(43), 17095–17107. 10.1523/JNEUROSCI.0890-13.2013

76. Cheng, H.-C. et al., (2010). Clinical progression in Parkinson disease and the neurobiology of axons. Ann of Neurol, 67(6), 715–725. 10.1002/ana.21995

77. Grünhage, F. et al., (2000). Systematic screening for DNA sequence variation in the coding region of the human dopamine transporter gene (DAT1). Mol Psychiatry, 5(3), 275–282. 10.1038/sj.mp.4000711

78. Drossos, C., Hunter, S.J. Juvenile Parkinson’s Disease. In: Kreutzer, J.S., DeLuca, J., Caplan, B., eds. Encyclopedia of Clinical Neuropsychology. Springer; (2011). https://doi-org.myaccess.library.utoronto.ca/10.1007/978-0-387-79948-3_1558

79. Klepac, N. et al., (2013). An update on the management of young-onset Parkinson’s disease. Degener Neurol Neuromuscul Dis, 2, 53–62. 10.2147/DNND.S34251

