## Supplemental Figures for "Characterization of a novel mouse model of Dopamine Transporter Deficiency Syndrome and pharmacological therapeutic strategies"

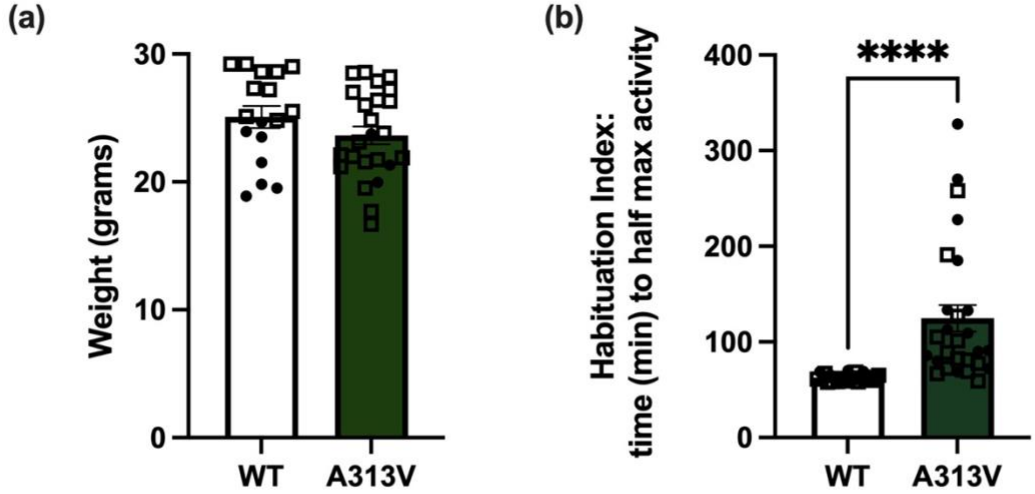


**Supplemental Figure 1. Comparison of Body Weight and Habituation Index in WT and A313V Mice.** (a) Weight in grams of wildtype (WT) (n=16) and dopamine transporter (DAT) A313V knock-in (A313V) (n=23) mice (25.08 ± 0.85vs 23.63 ± 0.70 grams, t=1.308, df= 39, p=0.1986) (n=17-24) (b) Habituation index (time in minutes to reach half of maximal activity) in the open field test in WT and A313Vmice. A313V mice took significantly greater time than WT mice to reach half max activity, suggestive of a decreased ability to behaviorally habituate to the novel environment (62.93 ± 0.72 vs 124.71 ± 13.75 minutes, t=4.402, df=51, p<0.0001) (n=26-27). Closed circular points represent values from female mice, and open square points represent values from male mice. Results are presented as mean ± SEM. Student's unpaired, two-tailed t-tests were conducted, ns p>0.05, ****p<0.0.0001

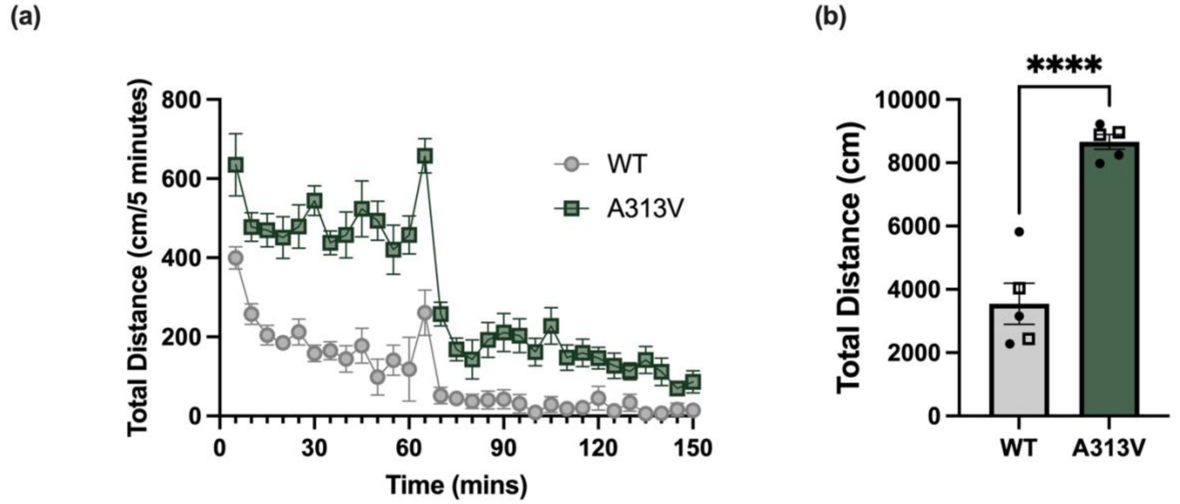


**Supplementary Figure 2. Baseline Locomotor Activity in 15-month old WT and A313V Mice.** Baseline locomotor behavior of wildtype (WT), and dopamine transporter (DAT) A313V knock-in (A313V). (a) Total distance was recorded in 5-minute bins for 150 minutes in an open field chamber. Animals were given i.p injections of saline at 60 minutes. A313V mice traveled significantly greater cumulative average levels of (b) total distance (3548.40 ± 648.59 vs 8659.40 ± 233.89 cm, t=7.413, df=8, p<0.0001) (n=5-6). Closed circular points represent values from female mice, and open square points represent values from male mice Results are presented as mean ± SEM. Student's unpaired, two-tailed t-tests were conducted, ***p≤0.001

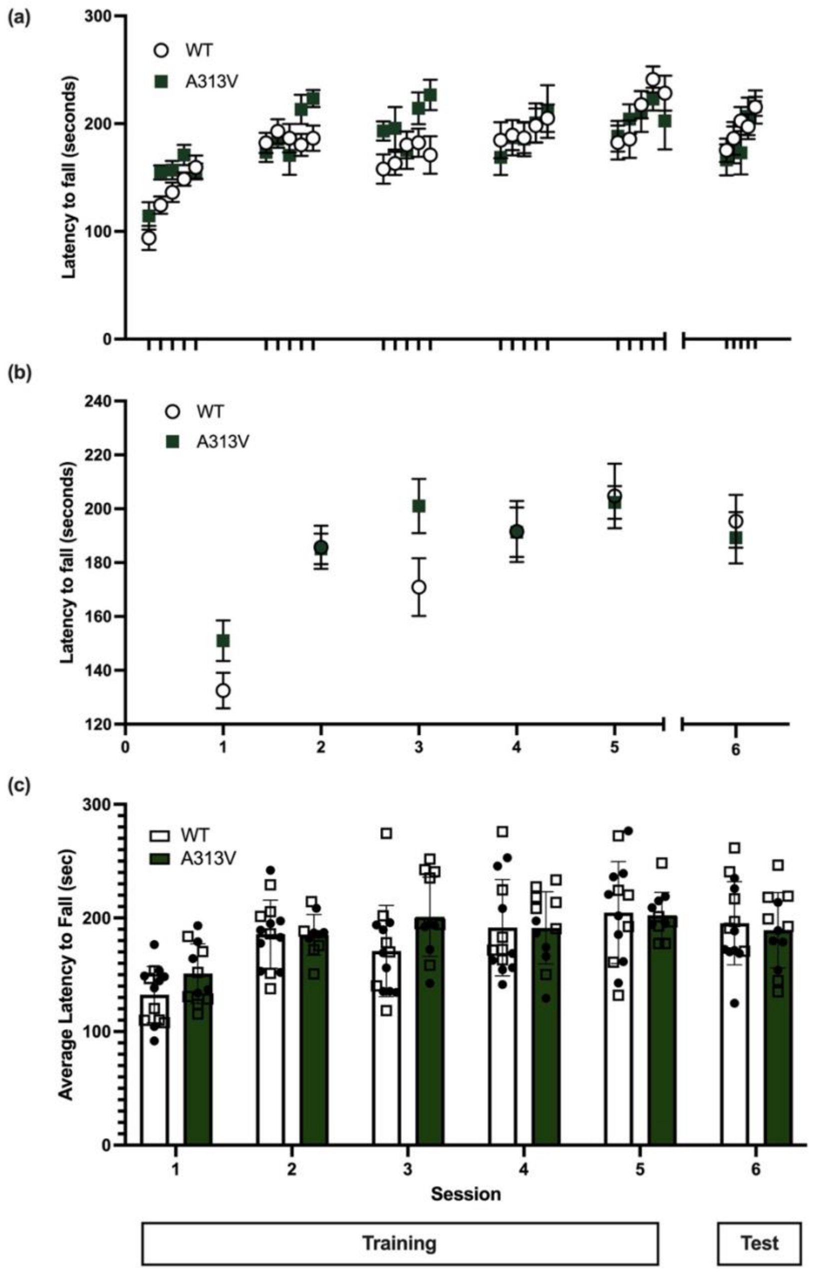


**Supplementary Figure 3. Motor Learning and Coordination in WT and A313V Mice.** Wildtype (WT) and dopamine transporter (DAT) A313V knock-in (A313V mice were trained on the rotarod for 5 sessions (Sessions 1-5). After a three day break, the mice were tested (Session 6). (a) Latency to fall in each trial. (b,c) Average latency to fall during each session. Results are presented as mean ± SEM. A mixed effect analysis was conducted. There was no significant effect of genotype ($F_{1,13}$=0.6625 p=0.4303), or a genotype by trial interaction ($F_{5,50}$=1.727 p=0.1455). There was a significant effect of trial ($F_{5.65}$ =14.79, p<0.0001) (n=11-14). Closed circular points represent values from female mice, and open square points represent values from male mice. N.s. p>0.05

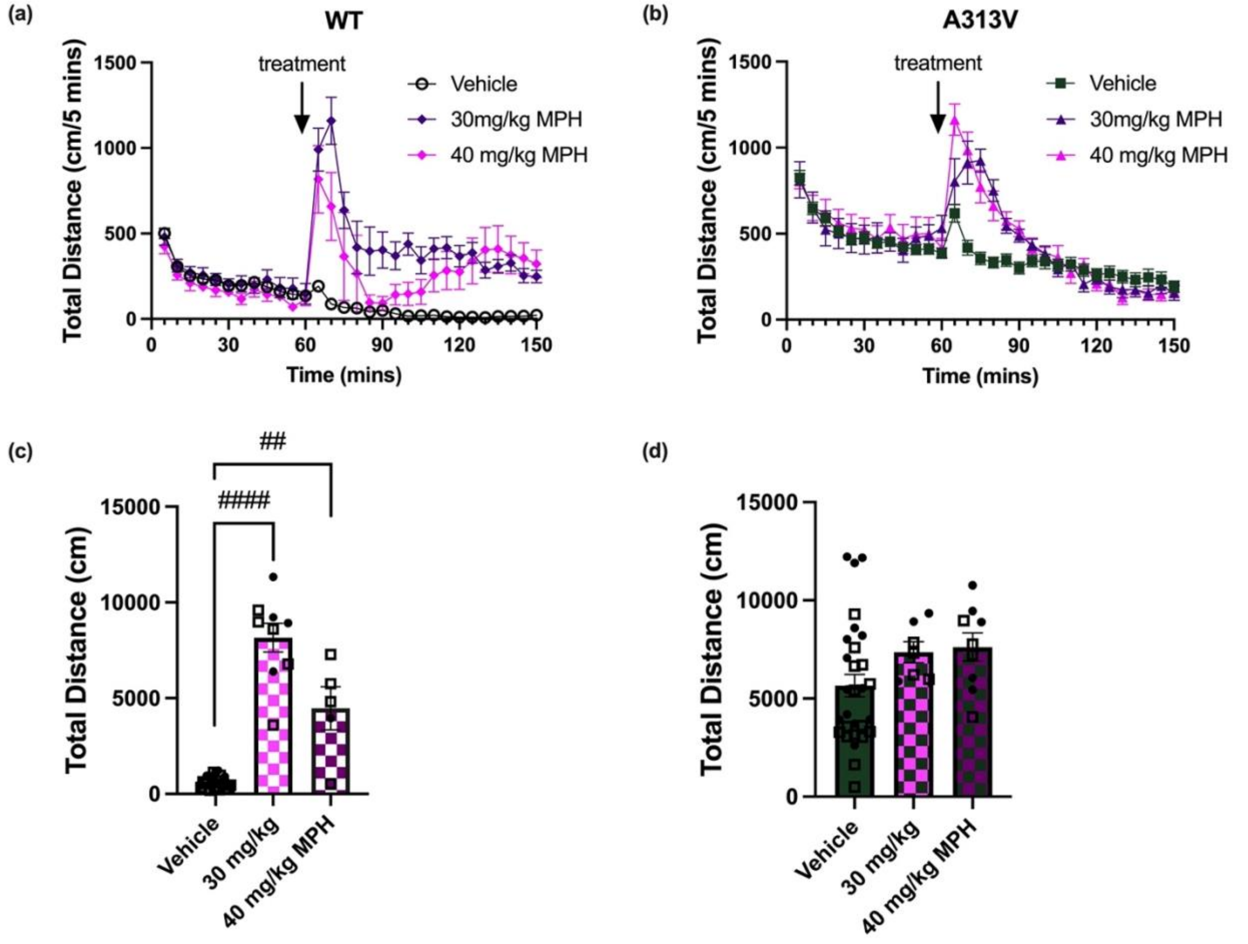


**Supp. Fig. 4. Effect of methylphenidate on locomotor activity.** Total distance was recorded in 5-minute bins in wildtype (WT) and dopamine transporter (DAT) A313V knock-in (A313V) mice over a period of 150 minutes. The test began with a 60 minute baseline followed by the intraperitoneal (i.p.) administration of 30 mg/kg or 40 mg/kg methylphenidate (MPH) at the 60-minute time point. Drug intervention is marked with an arrow. (a) total distance traveled in WT and (b) A313V mice treated with 30 mg/kg MPH or vehicle. (c) bar graph representing the sum of total distance traveled post-treatment of 30mg/kg MPH or vehicle in WT and (d) A313V mice (n=5-29). A two-way ANOVA showed a significant main effect of treatment ($F_{2,79}=30.25$, $p<0.0001$), genotype ($F_{1,79}=17.31$, $p<0.0001$), and a significant genotype x treatment interaction ($F_{2,79}=10.59$, $p<0.0001$). Tukey's multiple comparisons showed that, in WT mice, total distance traveled upon treatment of both 30 mg/kg and 40 mg/kg was increased compared to vehicle (652.62 ± 63.32 vs 8156.67 ± 750.17 cm, $p<0.0001$; 652.62 ± 63.32 vs 4468.40 ± 1127.82, $p=0.0020$). For A313V mice, there were no significant differences between vehicle and 30 mg/kg MPH (5658.48 ± 576.75 vs 7357.29 ± 535.62 cm, $p = 0.1693$), and vehicle and 40 mg/kg MPH (5658.48 ± 576.75 vs 7623.44 ± 715.31 cm, $p = 0.0581$). Closed circular points represent values from female mice, and open square points represent values from male mice. Results are presented as mean ± SEM. For post hoc effects: n.s. $p>0.05$, ##$p\leq0.01$, ####$p<0.0001$

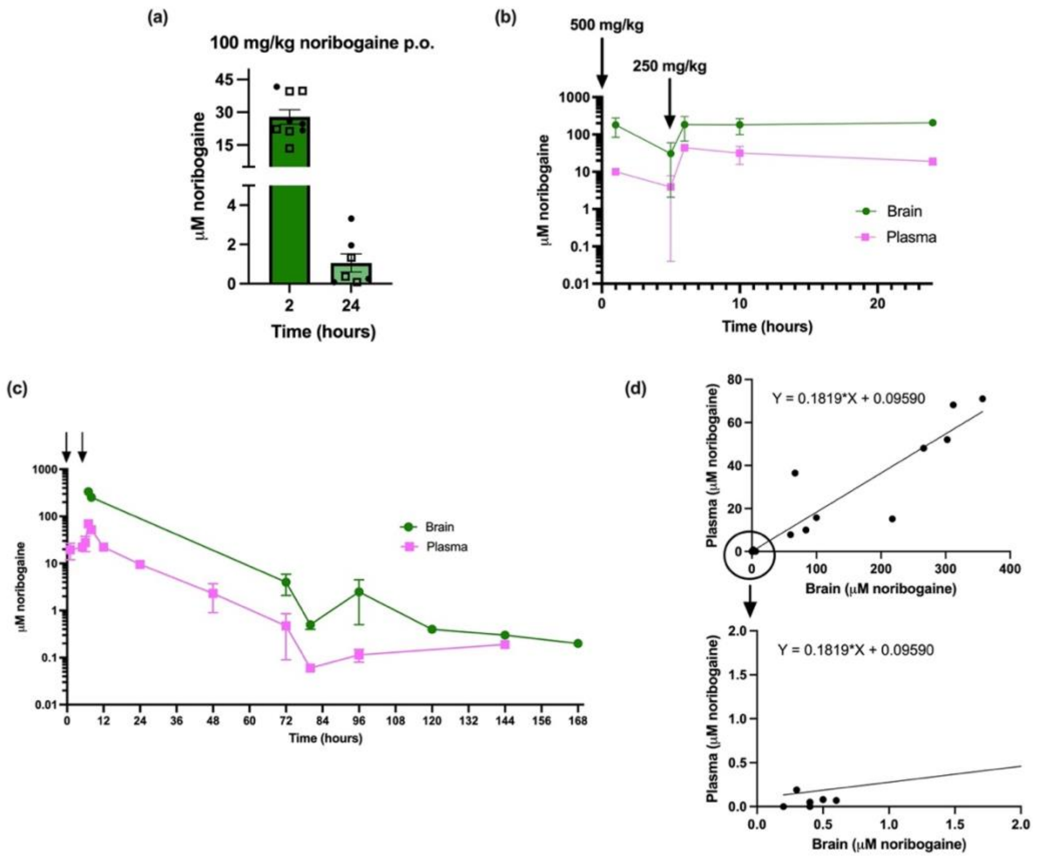


**Supplemental Figure 5. Establishment of noribogaine dosing regimen.** (a) Noribogaine concentrations (μM) in mouse cerebellar brain tissue at 2 and 24 hours following a single oral gavage dose of 100 mg/kg noribogaine (n=7-9). (b) Noribogaine concentrations (μM) in cerebellar (brain) tissue over a 0-30 hour time course following a dose of 500 mg/kg oral gavage at hour 0 and 250 mg/kg oral gavage at hour 5. Arrows indicate the timing and dose of noribogaine administration. Animals whose tissue was collected at the 5 hour time point did not receive the second 250 mg/kg administration. Brains and plasma were collected from 2 animals per timepoint, with the exception of hour 1, where plasma was collected from one animal only (c) Time course of noribogaine concentrations (μM) in cerebellar (brain) and plasma samples from 0 to 168 hours post-gavage of noribogaine dosing regimen to show extended pharmacokinetics of noribogaine in both tissues. Arrows indicate time of noribogaine administration. Tissue was collected from two animals per timepoint, with the exception of hour 5, hour 120, hour 144, and hour 168. (d) Relationship between cerebellar (brain) noribogaine (μM) and plasma noribogaine (μM) concentrations from (b,c), illustrating correlated distribution between compartments at multiple timepoints post-administration. The lower panel is a close up of the upper panel around x,y=2,2. Closed circular points represent values from female mice, and open square points represent values from male mice. Data are presented as mean ± SEM. The y-axis in graphs (a) and (d) is presented on a linear scale, while the y-axis in graphs (b) and (c) is presented on a logarithmic scale. Results are presented as mean ± SEM.

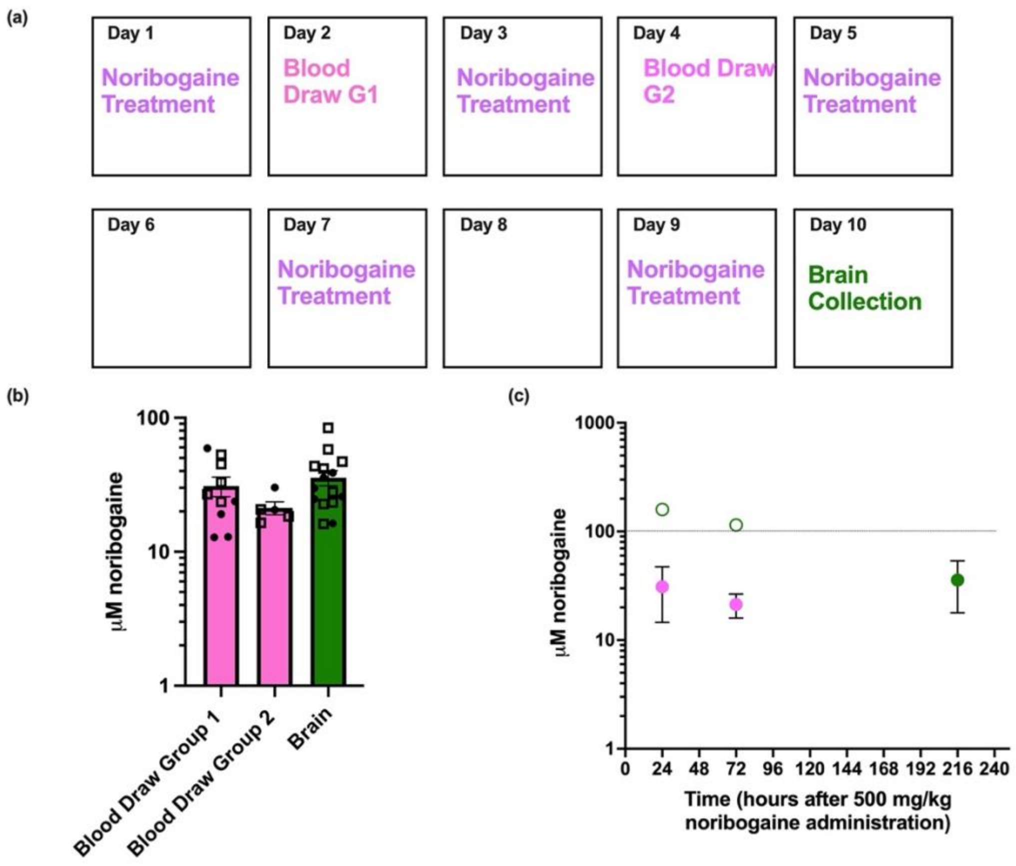


**Supplemental Figure 5. Noribogaine concentrations in the brain and blood following chronic dosing.** (a) Experimental timeline for chronic noribogaine administration. Noribogaine was administered ("Noribogaine Treatment") via oral gavage according to the established regimen, at 500 mg/kg+ 250 mg/kg five on days 1, 3, 5, 7, and 9, with dosing occurring every 48 hours. Blood samples were collected 24 hours post-administration on day 2 from Group 1 animals and on day 4 from Group 2 animals (n=5-10). Brain samples (cerebellar tissue) were collected from all animals (n= 15) on day 10, 24 hours after the final noribogaine administration on day 9. (b) Noribogaine concentrations measured in blood samples from Group 1 and Group 2 and in brain (cerebellar tissue) samples from all animals collected on day 10. Concentrations are reported in micromolar (μM). (c) Scatter plot illustrating the relationship between plasma noribogaine levels and predicted brain concentrations. Pink circles indicate plasma noribogaine concentrations measured 24 hours after oral administration of noribogaine (500 mg/kg followed by 250 mg/kg) on days 1 and 3, with values taken at hours 24 and 72 post-dosing. Green-outlined circles represent the predicted brain concentrations calculated from plasma levels using the equation Y= 0.1819 *X + 0.0950 derived in Supplementary Figure 5*d*, where Y is equal to the plasma noribogaine concentration in μM and X is equal to the brain noribogaine concentration in μM. Predicted brain noribogaine concentrations based on mean plasma levels of 30.931 μM at 24 hours and 21.234 μM at 72 hours are 169.52 μM and 116.21 μM, respectively. The filled green circle represents the actual mean brain noribogaine concentration measured at hour 216, which was 35.7 μM. Closed circular points represent values from female mice, and open square points represent values from male mice. The y-axis in (b,c) are presented on a logarithmic scale. Results are presented as mean ± SEM.

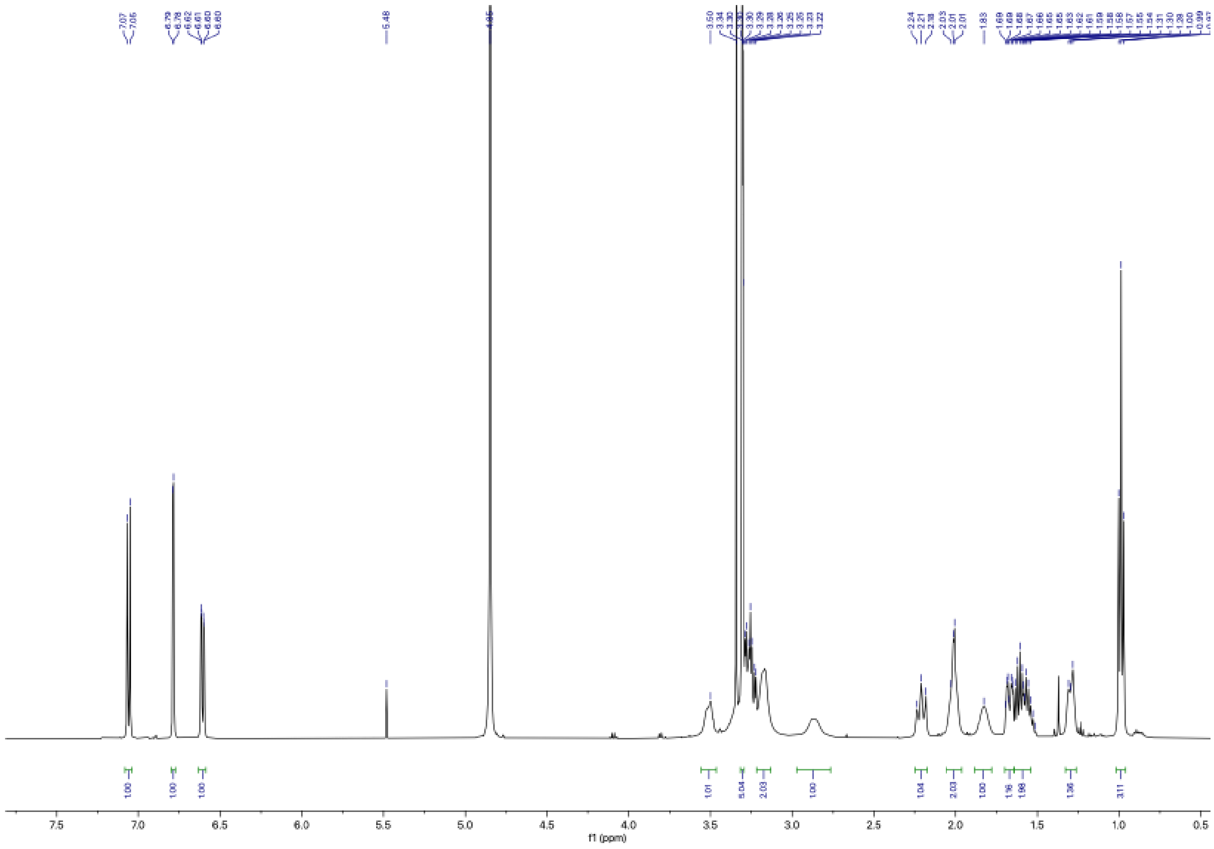

**Supplemental Figure 6. 500 MHz $^{1}$H NMR of noribogaine HCl in $CD_3OD$.**

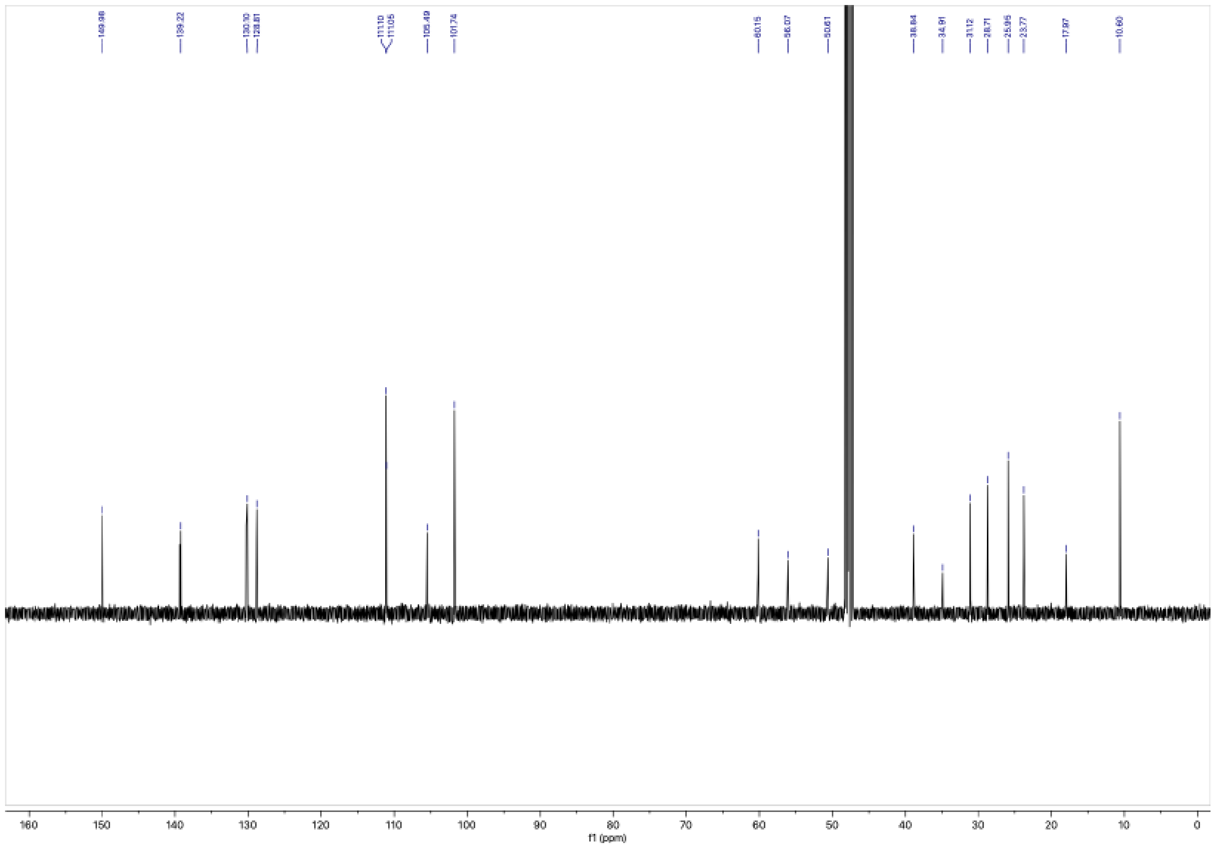


**Supplemental Figure 7. 125 MHz $^{13}C$ NMR of noribogaine HCl in $CD_3OD$.**
